# Gβγ dually regulates the M-current via increased channel surface expression and PIP2 sensitivity

**DOI:** 10.64898/2026.09.15.751787

**Authors:** Boris Shalomov, Iuliia Papa-Dmitrieva, Tal Keren Raifman, Sharon Weiss, Flavia De Martino, Adi Gali-Sayag, Joel A. Hirsch, Maurizio Taglialatela, Vincenzo Barrese, Kerstin Zühlke, Enno Klussmann, Iain Greenwood, Ilana Lotan, Nathan Dascal

## Abstract

The M-current, generated by voltage gated K_V_7.2/7.3 channels, sets the threshold for neuronal action potential and acts as a key brake on repetitive firing. The M-current is tightly regulated by signaling molecules, such as calmodulin and phosphatidylinositol-4,5-bisphosphate (PIP_2_). Here, we show that coexpression of the ubiquitous subunit dimer of heterotrimeric G-proteins, Gβγ, with K_V_7.2/7.3 in *Xenopus laevis* oocytes doubles maximal M-current. This regulation requires free, prenylated, membrane–associated Gβγ and operates via two distinct mechanisms: 1) increasing plasma membrane (PM) channel density and 2) stabilizing K_V_7.2/7.3-PIP₂ coupling. Conversely, Gβγ scavengers reduce basal K_V_7.2/7.3 current and weaken PIP_2_ coupling. Proximity ligation assays confirm colocalization of Gβγ and K_V_7.2/7.3 in the PM. Peptide array and AlphaFold modeling identify putative interaction sites on the channel’s cytoplasmic domain. Finally, the disease–causing Gβ_1_ variant I80N abolished Gβγ–induced M-current potentiation. Together, these findings establish Gβγ as a significant physiological regulator and potential site of vulnerability in neuronal M-current function.

## Introduction

Gβγ – the ubiquitous, obligatory heterodimer of Gβ and Gγ subunits of heterotrimeric G proteins – regulates plethora of cellular processes (*1, 2*). Canonically, most Gβγ–induced regulations are initiated following its release from the heterotrimeric G protein, Gαβγ, caused by agonist activation of G-protein coupled receptors (GPCRs) (*3*); however, some regulations of specific cellular targets by Gβγ are GPCR–independent (*1, 4, 5*).

Gβγ directly engages a broad set of effectors, activating G-protein–gated inwardly rectifying K^+^ channels (GIRK/Kir3), inhibiting voltage–gated Ca^2+^ (Ca_v_2) channels and presynaptic SNARE proteins, and regulating adenylyl cyclases, phospholipases, and protein kinases, with the functional outcome depending on the specific Gβ and Gγ isoforms and on prenylation of the Gγ subunit that anchors the Gβγ dimer to the membrane (*2, 6, 7*). In the nervous system, the prevailing effect of free Gβγ (basal or released following activation of G_i/o_ proteins) is inhibition of excitability, neuronal activity and neurotransmitter release due to the inhibition of Ca_V_2 channels and SNARE proteins and activation of GIRK (*8–11*). GIRK is a paradigmatic effector of Gβγ, activated by direct Gβγ binding through a membrane–delimited mechanism. Its function requires phosphatidylinositol-4,5-bisphosphate (PIP₂), with Gβγ acting cooperatively by strengthening the channel–PIP_2_ interaction (*12–15*).

More recently it has been discovered that Gβγ augments the activity of another K^+^ channel, the voltage–gated K_V_7.4 (*16*), one of the five members of the K_V_7 family of voltage– gated K^+^ channels encoded by *KCNQ1-5* genes. K_V_7 channels generate slowly activating, non-inactivating currents active near the resting membrane potential that set the threshold for excitation in many cell types (*17, 18*). Like GIRKs, K_V_7 channels are also reliant upon direct binding of PIP2, which secures coupling of voltage sensor movement to pore opening through binding at multiple sites, including the S2-S3 and S4-S5 linkers and the proximal C-terminus (*19–22*). The latter is also constitutively associated with calmodulin, which is required for channel assembly and Ca^2+^-dependent regulation (*23, 24*). Gβγ augmentation of K_V_7.4 is mediated by an increase in voltage sensitivity and activation rate that synergizes with PIP_2_ action (*16, 25*). However, effects of Gβγ on other K_V_7 isoforms are unknown.

In neurons, proteins encoded by *KCNQ2* and *KCNQ3* (K_V_7.2/7.3) are the principal molecular correlate of the neuronal M-current (*26, 27*), a potent regulator of neuronal activity (*28, 29*). K_V_7.2 and K_V_7.3 concentrate at axon initial segments and nodes of Ranvier, where they act as a powerful brake on repetitive firing (*30, 31*). Loss-of-function (LoF) mutations in either subunit cause benign familial neonatal epilepsy or, in severe cases, developmental and epileptic encephalopathy (*32*). Mutations in *GNB1*, the gene encoding the Gβ_1_ subunit also cause developmental encephalopathy with affected individuals presenting with global developmental delay, hypotonia or dystonia, and epilepsy (*33, 34*). Functional studies have tied specific variants to altered GPCR–G protein and Gβγ–effector coupling, including a gain-of-function toward GIRK channels that drives seizures in a K78R mouse model (*35*), and loss-of-function effects for other substitutions (*36–38*).

This study investigates whether K_V_7.2/7.3 channels are modulated by Gβγ and if this regulation is altered by GNB1 encephalopathy–linked mutants. We show that Gβγ interacts with multiple intracellular regions of K_V_7.2 and K_V_7.3, colocalizes with both subunits, increases whole-cell K_V_7.2/7.3 currents, and strengthens functional coupling between PIP_2_ and the K_V_7.2/7.3 channel in heterologous expression systems. *GNB1* encephalopathy–associated variants differentially alter Gβγ regulation of the M-current, offering a potential mechanistic link to neurodevelopmental disease.

## Results

### Gβγ enhances the K_V_7.2/7.3 M-current without significantly altering voltage dependence

To test whether Gβγ modulates the M-current, we coexpressed K_V_7.2 and K_V_7.3 channel subunits with or without Gβ_1_γ_2_ (Gβγ) in *Xenopus laevis* oocytes, by using cRNA doses established to robustly activate GIRK channels (*39*). We recorded whole-cell currents by two-electrode voltage clamp (TEVC) and found that coexpression of Gβγ produced a substantial increase in current amplitude across the activation range (Fig. 1A, B). The current–voltage (I-V) relationship was fitted with Boltzmann I-V equation. The maximal conductance (G_max_) was approximately doubled by Gβγ (Fig. 1C, P < 0.0001), establishing that Gβγ is a strong positive modulator of K_V_7.2/7.3. Across oocyte batches, Gβγ–mediated enhancement ranged from 1.4-to ∼3-fold (Table S1).

**Fig. 1.**
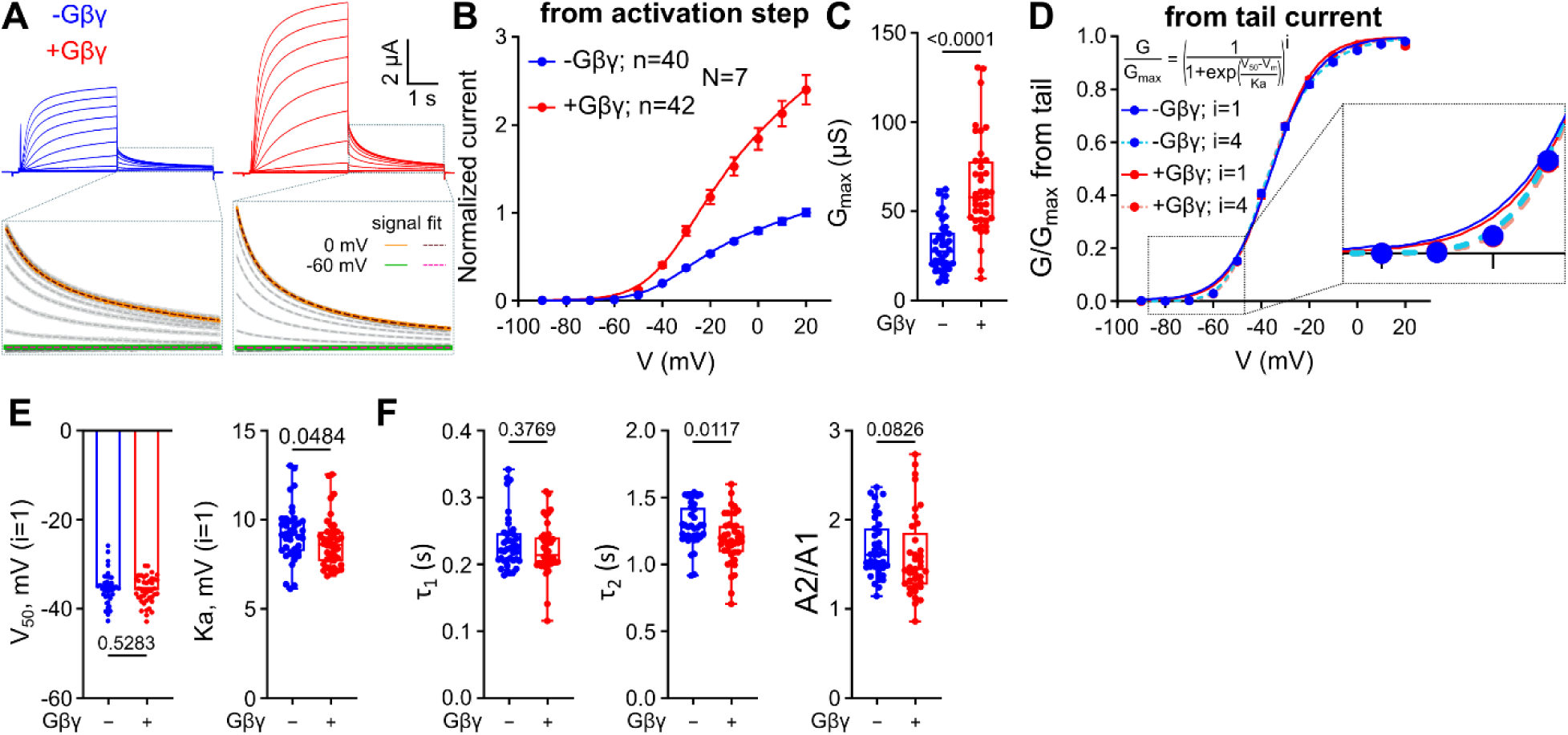
Gβ_1_γ_2_ robustly enhances the K_V_7.2/7.3 M-current with little change in voltage-dependence of gating. K_V_7.2 and K_V_7.3 were co-expressed in *Xenopus laevis* oocytes without (−Gβγ) or with (+Gβγ) Gβ_1_γ_2_, and currents were recorded by two-electrode voltage clamp using protocols detailed in Methods. Panels A-F show results from the same set of experiments. The number of oocytes (n) and experiments (N) were: -Gβγ, n = 40; +Gβγ, n = 42; N = 7. (**A**) Representative current recordings in the absence (blue) and presence (red) of Gβγ. Currents were elicited by steps from -90 to increasing voltages in 10 mV steps, followed by post-test step to -60 mV. Insets show tail-current relaxations at -60 mV with bi-exponential fits. (**B**) Mean normalized current-voltage (I-V) relationships from the activation steps. (**C**) Maximal conductance (G_max_); Gβγ approximately doubled G_max_ (p < 0.0001). (**D**) Mean G-V relationships from tail current fit to a Boltzmann function raised to the first (i = 1) or fourth (i = 4) power (equation inset). The boxed near-threshold region is enlarged at right and shows a visually slightly better, but not statistically significant, fit with i=4. (**E**) Half-activation voltage (V_50_) and slope factor (K_a_) for i = 1 (E). (For i = 4, see Fig. S1G). (**F**) Coexpression of Gβγ does not appreciably affect deactivation kinetics: fast (τ1) and slow (τ2) time constants and relative amplitudes of the two relaxation components (A2/A1).

Notably, this current potentiation occurred without substantial shifts in the voltage dependence of activation, as determined by tail current (I_tail_) analysis (see Methods). Conductance–voltage (G-V) relationships were virtually superimposable in the presence and absence of Gβγ and were well described both by a single Boltzmann function (i = 1), which treats activation as one concerted transition, and by a four-power Boltzmann function (i = 4), the classic Hodgkin-Huxley n⁴ formalism (*40, 41*) in which four independent voltage sensing domains must engage before the pore opens (Fig. 1D). Although individual batches showed a subtle trend toward more hyperpolarized half-activation voltages (V_50_), on average mean values for V_50_ and the slope factor K_a_ were not significantly altered by coexpression of Gβγ (Fig. 1E, Fig. S1G). Analysis of deactivation kinetics revealed no change in the fast time constant (τ_1_) and only a small change in the slow time constant (τ_2_), with the relative amplitude of the two components (A2/A1) unchanged (P = 0.0826; Fig. 1F). These data indicate that coexpressed Gβγ augments the M-current without major alterations in the voltage dependence of gating.

We observed similar effects for homotetrameric K_V_7.2 channels (Fig. S1). Gβγ significantly increased the normalized current at 0 mV for heteromeric K_V_7.2/7.3 (p = 0.0024) and for homomeric K_V_7.2 (p = 0.0024). There was also a tendency of Gβγ to increase current via homomeric K_V_7.3 channels, but it did not reach statistical significance (p = 0.0646) (Fig. S1, A-D). With all subunit configurations, V_50_ and K_a_ were not significantly affected by Gβγ (Fig. S1E, F).

### Gβγ elevates the surface expression of K_V_7.2/7.3 channels

The increase in Gmax without significant changes in voltage dependence suggested that Gβγ may increase channel density at the plasma membrane. To test this directly, we prepared giant plasma membrane patches (GMPs) isolated from *Xenopus* oocytes (*42*). In this configuration, outer membrane surfaces adhere to the coverslip while cytosolic surfaces face the bath solution, enabling quantitative confocal imaging of surface-expressed proteins labeled with antibodies targeting intracellular epitopes (Fig. 2). Coexpression of Gβγ significantly increased the plasma membrane (PM) density of K_V_7.2/7.3, as determined by quantifying the fluorescence of YFP-tagged K_V_7.3 subunit (Fig. 2A, B). Immunostaining with a Gβ_1_ antibody gave a faint signal for the endogenous oocyte Gβ, but much stronger signals in Gβ_1_–expressing oocytes, confirming the robust accumulation of the expressed Gβγ at the PM (Fig. 2A, C). In parallel, in the same set of experiments, the whole-cell currents recorded at 0 mV were also increased (Fig. 2D).

**Fig. 2.**
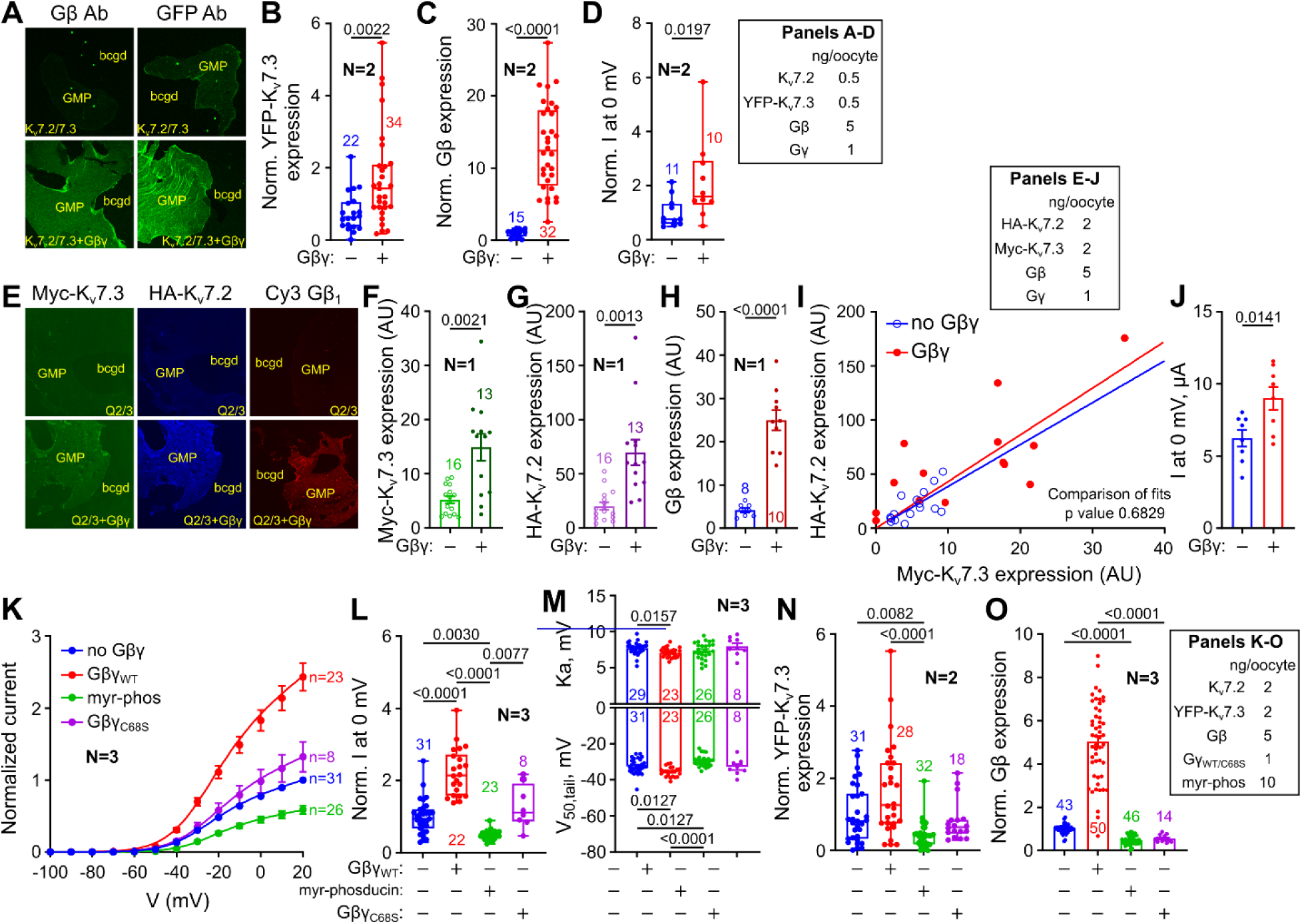
Gβγ elevates the plasma membrane abundance of K_V_7.2/7.3 channels and requires prenylated Gβγ. (**A**) Confocal images of GMPs from oocytes expressing K_V_7.2/YFP-K_V_7.3 ± Gβγ, stained with antibodies for Gβ and YFP. Coexpression of Gβγ increased surface YFP-K_V_7.3 expression (**B**), surface Gβ_1_ (**C**), and M-current (**D**; normalized current at 0 mV is shown). Amounts of injected RNAs in A-D are shown in the inset in (D). (**E-J**), Gβγ increases surface abundance of both channel subunits oocytes expressing HA-K_V_7.2/Myc-K_V_7.3 ± Gβγ. Amounts of RNA are shown in the inset in (I). (**E**) Confocal images of GMPs double-labeled by antibodies for HA and Myc, and Gβ_1_. (**F**, **G**) Expression of Gβγ increases surface levels of Myc-K_V_7.3 and HA-K_V_7.2. (**H)** Gβ_1_ is robustly expressed in PM. (**I**) Per-GMP correlation of surface HA-K_V_7.2 versus Myc-K_V_7.3 without and with Gβγ. The slopes of linear regressions were almost identical (extra sum-of-squares F test; p = 0.6829). (**J**) Gβγ increases the M-current in oocytes of the same experiment. (**K-O**) Effects of coexpression of Gβ_WT_γ, myr-phosducin (myr-phos), or Gβγ_C68S_ on KV7.2/YP-K_V_7.3. Amounts of injected RNAs are shown in the inset in (O). (**K)** Mean normalized I-V relationships. RNA (ng/oocyte): HA-K_V_7.2:YFP-K_V_7.3 2:2, Gβ_WT_:Gγ 5:1, Gβ_C68S_:Gγ 5:1, myr-phosducin 10. (**L**) Normalized currents at 0 mV. (**M**) Comparison of V_50_ and K_a_ between treatments. (**N**) Surface YFP-K_V_7.3 expression. (**O**) Coexpression of K_V_7.2/7.3 + myr-phosducin decreased endogenous Gβγ levels in PM compared to K_V_7.2/7.3 alone (Welch’s t test; p<0.0001) and coexpressed Gβγ_C68S_ significantly decreased Gβγ levels in PM compared to Gβ_WT_γ (Mann-Whitney test; p<0.0001).

Next, we generated N-terminally tagged HA-K_V_7.2 and Myc-K_V_7.3 to determine whether the relative abundance of these subunits at the PM changed with Gβγ coexpression. Coverslips with GMPs expressing HA-K_V_7.2/Myc-K_V_7.3, with or without Gβγ, were double–labeled with antibodies against HA and Myc; a separate set of coverslips was also labeled with an antibody against Gβ1 (Fig. 2E). HA and Myc labeling confirmed the presence of both K_V_7.2 and K_V_7.3 at the PM. The PM levels of both HA-K_V_7.2 and Myc-K_V_7.3 increased in the presence of Gβγ, as did the current measured at 0 mV in oocytes from the same batch (Fig. 2E-H, J). The surface levels of K_V_7.2 and K_V_7.3 were tightly correlated, and the slope of this relationship was indistinguishable in the presence and absence of Gβγ (comparison of linear fits, P = 0.6829; Fig. 2I). These results indicate that Gβγ increases the PM abundance of the two subunits in a coordinated manner rather than selectively promoting trafficking of one subunit.

To determine whether the increase in K_V_7.2/7.3 surface expression requires free, membrane–associated Gβγ, we used two complementary approaches. First, we coexpressed Gβ with GγC68S, a prenylation–deficient mutant carrying a Cys68Ser substitution that prevents Gβγ membrane anchoring while preserving whole-cell expression of the Gβγ_C68S_ protein (*39*). While wildtype Gβγ (Gβ_WT_γ) strongly increased both whole-cell K_V_7.2/7.3 currents and surface channel levels (Fig. 2K, L), Gβγ_C68S_ failed to increase both currents (Fig. 2K,L) and K_V_7.2/7.3 surface expression in GMPs (Fig. 2N; p>0.99 vs K_V_7.2/7.3 alone), suggesting that membrane attachment of Gβγ is required for both effects. Second, we coexpressed K_V_7.2/7.3 with myr-phosducin, a membrane–targeted scavenger that binds and sequesters free Gβγ and greatly decreases the basal current of GIRK1/2, the archetypal Gβγ–activated K^+^ channel, which is largely Gβγ– dependent (*43*). Myr-phosducin strongly reduced the basal K_V_7.2/7.3 current (Fig. 2K,L, Fig. S2B, E), similar to GIRK1/2 used here as control (Fig. S2A).

Incidentally, in this set of experiments (N=3), the small hyperpolarizing change in V_50_ caused by coexpression of Gβγ reached statistical significance (∼3 mV, p=0.006), whereas Gβγ_C68S_ did not change V_50_ (Fig. 2M). Strikingly, myr-phosducin produced a small but significant shift in the opposite direction, increasing V_50_ by ∼3 mV compared with oocytes expressing K_V_7.2/7.3 alone (P = 0.0175; Fig. 2M, Fig. S2). Thus, V_50_ differed by ∼6 mV between channels expressed with Gβγ and those in which Gβγ was scavenged by phosducin (P < 0.0001; Fig. 2M). K_a_ was slightly reduced by Gβγ but not consistently affected by phosducin or Gβγ_C68S_ (Fig. 2M). We also measured channel surface expression using K_V_7.2/YFP-K_V_7.3 in two experiments in this series. Gβγ increased surface expression by 67%, although this did not reach statistical significance, whereas myr-phosducin reduced it by ∼60% (P = 0.0082); Gβγ_C68S_ had no effect (Fig. 2N). Collectively, the mild but consistent, opposite changes in V_50_ produced by Gβγ addition *versus* its removal by scavenging suggest that Gβγ may regulate K_V_7.2/7.3 gating in addition to increasing surface expression. Moreover, the robust reduction in surface expression caused by phosducin suggests that Gβγ is an important cofactor for K_V_7.2/7.3 surface expression in this heterologous system.

### Gβγ stabilizes functional K_V_7.2/7.3–PIP₂ coupling

We next sought to determine whether Gβγ affects K_V_7.2/7.3-PIP₂ coupling. To this end, we coexpressed K_V_7.2/7.3, with or without Gβγ, together with *Ciona intestinalis* voltage–sensitive phosphatase (Ci-VSP), which cleaves PIP₂ (*44*). Ci-VSP was activated by stepping the voltage to 20 mV for 15 s, resulting in PIP₂ cleavage and consequent depletion in the PM. Accordingly, K_V_7.2/7.3 channel activity decayed over time; notably, the decay appeared slower in the presence of Gβγ (Fig. 3a, Fig. S3A).

**Fig. 3.**
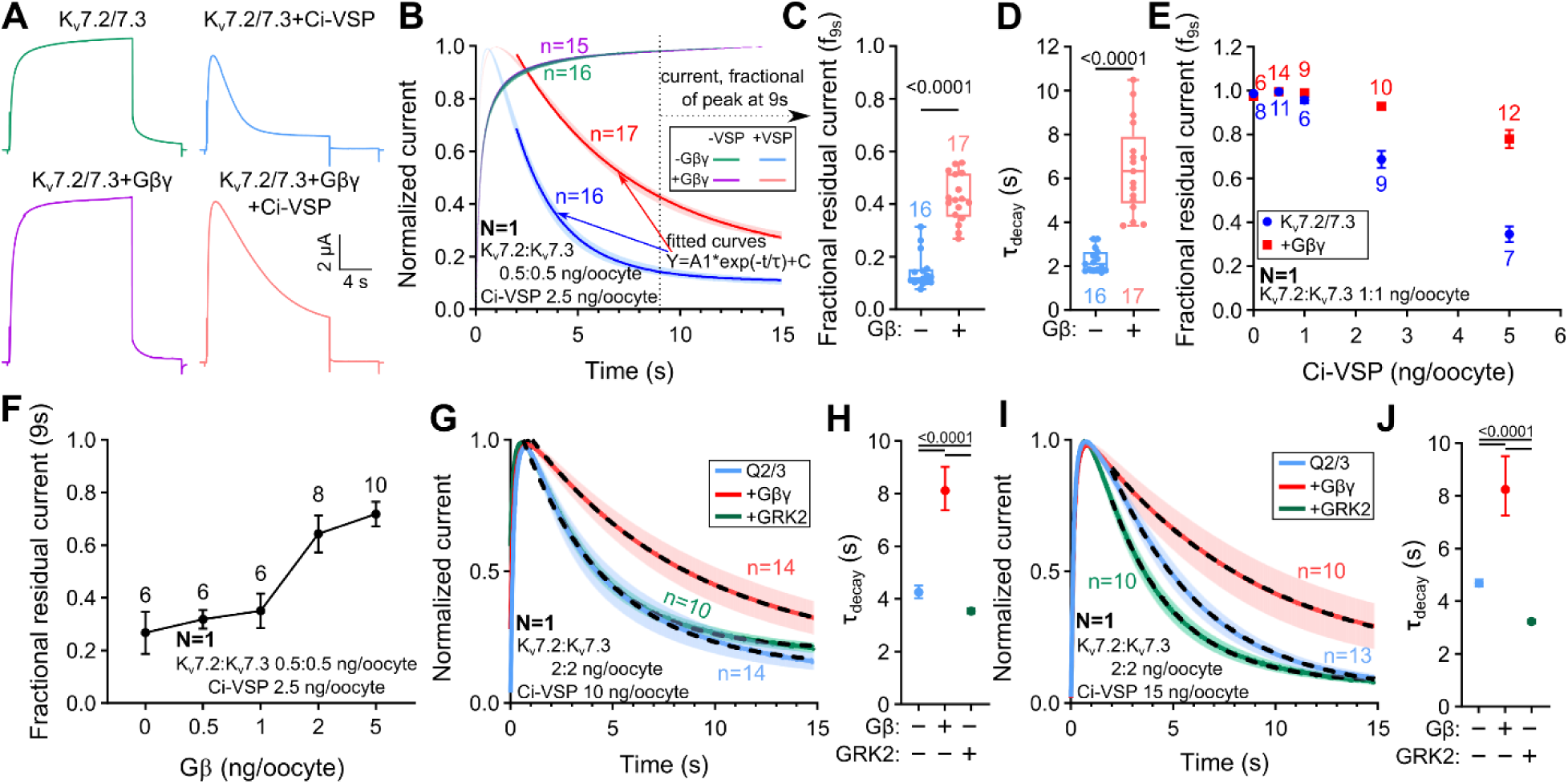
Gβγ stabilizes functional KV7.2/7.3-PIP_2_ coupling. Currents were elicited by a 15 s voltage step to +20 mV, to both trigger the M-current and activate Ci-VSP. (**A-D**) Gβγ slows down M-current decay caused by the activation of Ci-VSP. RNA doses are shown in (B). (**A**) Representative current records for K_V_7.2/7.3 expressed alone or with Gβγ, with or without Ci-VSP. (**B**) Averaged normalized current traces (mean±SEM) from a representative experiment. The dashed vertical line marks the 9-s quantification point. The decay phases were fit to a single-exponential function. (**C**) Gβγ increased the fraction of peak current remaining at 9 s. (**D**) Gβγ increases the τ_decay_ (derived from the exponential fits). (**E**) Gβγ slows down current decay in a range of Ci-VSP RNA doses. RNAs of K_V_7.2 and K_V_7.3 were 1 ng/oocyte for each subunit; Gβ:Gγ RNAs were 5:1 ng/oocyte. (**F**) Titrating Gβ RNA progressively increased the fraction of current resistant to PIP_2_ depletion. Gγ RNA was 1/5 of that of Gβ. (**G-J**) Coexpression of GRK2 accelerates M-current decay. Results of two separate experiments performed with the same batch of oocytes. The channel RNA does was 2 ng/oocyte of each subunit. The amount of GRK2 RNA was 10 ng/oocyte in the experiment of (G) and (H) and 15 ng/oocyte in the experiment of (I) and (J). Exponential fits have been performed on currents averaged from all cells in each group. The normalized currents in (G) and (I) are shown as mean±CI (95% confidence interval of τ from fit). (H) and (J) compare τ_decay_ (shown as mean±SEM) for different treatments, with 10 or 15 ng/oocytes of GRK2 RNA, respectively. Statistical tests: t-test between pairs of τ values with Bonferroni correction.

To quantify the effect of PIP_2_ depletion, we calculated the fractional residual current at 9 s relative to peak (f_9s_), when the difference between various conditions was usually well pronounced (Fig. 3A-C). As a second measure, we estimated the decay time constant (τ_decay)_ from single-exponential fit of current decay in each cell (Fig. 3A, B) or of the average current (Fig. 3G-J). Unlike f_9s,_ this analysis was not applicable in all cells, for example when current decay was too slow for a satisfactory exponential fit during the 15 s; in these cases only f_9s w_as calculated.

To optimize the assay, we titrated Ci-VSP cRNA while maintaining a constant amount of K_V7_.2/7.3 RNA (1 ng each subunit per oocyte). Ci-VSP–evoked current decay was maximal at 5 ng Ci-VSP cRNA per oocyte, establishing a 1:5 K_V_7.2/7.3:Ci-VSP cRNA ratio for robust detection of PIP_2_-depletion–induced decay (Fig. S3A,B). At this fixed ratio, increasing amounts of Gβγ progressively attenuated current decay, as reflected by an increase in f_9s_ and τ_decay_ values (Fig. 3C-F, Fig. S3C). Accordingly, with the same optimized 1:5 ratio used in the experiments of Fig. 3A–E (0.5 ng K_V_7.2 + 0.5 ng K_V_7.3 and 2.5 ng Ci-VSP RNA per oocyte), Gβγ coexpression significantly slowed Ci-VSP–induced K_V_7.2/7.3 current decay. These data indicate that Gβγ stabilizes functional coupling of K_V_7.2/7.3 channels to PIP_2_.

We next investigated whether reducing the availability of endogenous Gβγ would weaken functional K_V_7.2/7.3–PIP_2_ coupling. To this end, we coexpressed G protein–coupled receptor kinase 2 (GRK2), a high–affinity Gβγ–binding protein (*45*). Coexpression of GRK2 with either GIRK1/2 or K_V_7.2/7.3 significantly decreased the respective currents (Fig. S4), consistent with sequestration of endogenous free Gβγ. At K_V_7.2/7.3:Ci-VSP RNA ratios of 2:10 and 2:15, GRK2 coexpression accelerated Ci-VSP–induced current decay relative to K_V_7.2/7.3 alone, whereas Gβγ coexpression significantly attenuated the decay (Fig. 3G-J). Together, these reciprocal effects support a model in which endogenous Gβγ promotes functional K_V_7.2/7.3– PIP_2_ coupling, whereas its sequestration by GRK2 weakens this coupling.

### A proximity ligation assay identifies colocalization of K_V_7.2/7.3 subunits and Gβγ

Proximity ligation assay (PLA) detects proteins in close proximity within 40 nm, and is therefore suggestive of their colocalization or association *in situ* (*46*). To examine whether K_V_7.2/7.3 channels are in close proximity to Gβγ, we performed PLA in HEK293 (HEK) cells transfected with plasmids encoding K_V_7.2 and K_V_7.3, either individually or together, using antibodies directed against K_V_7.2 or K_V_7.3 and the Gβ subunit.

As shown in Fig. 4A, red PLA puncta, indicating proximity between K_V_7.2 and Gβ, were detected in HEK293 cells expressing K_V_7.2 alone (approximately 20 puncta/cell) and in cells co-expressing K_V_7.2 and K_V_7.3. By contrast, cells expressing K_V_7.3 alone displayed substantially fewer puncta (approximately 4 per cell), affirming specificity of the K_V_7.2 antibody (Fig. 4B). Similar results were obtained in assays of K_V_7.3–Gβ proximity. Cells expressing K_V_7.3 alone or co-expressing K_V_7.2 and K_V_7.3 exhibited abundant K_V_7.3–Gβ PLA puncta (approximately 30 and 40 puncta/cell, respectively), whereas cells expressing K_V_7.2 alone showed markedly fewer puncta (approximately 6 puncta/cell; Fig. 4C).

**Fig. 4.**
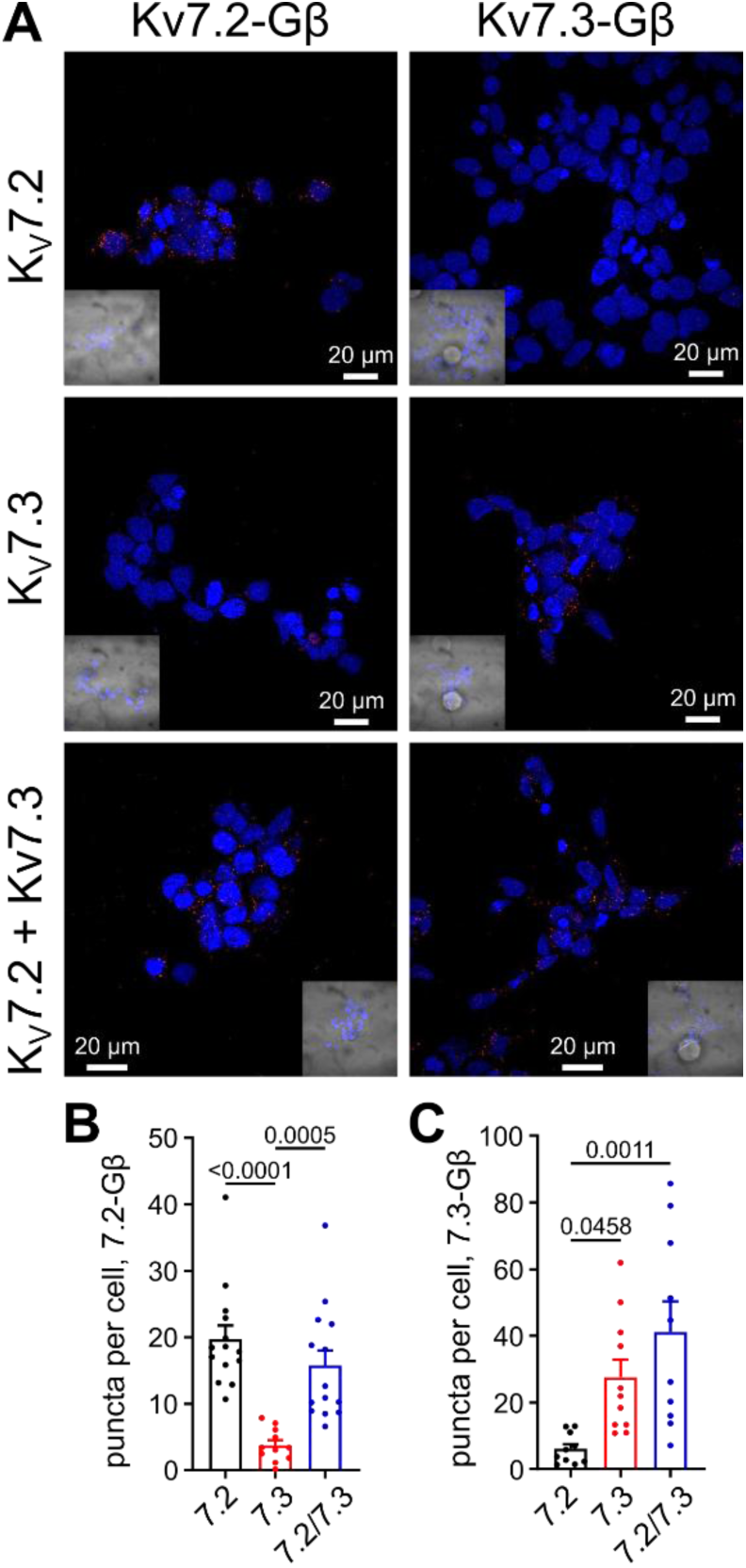
Proximity ligation assays identify direct interactions of KV7.2 and K_V_7.3 with endogenous Gβ. (**A**) Representative images of PLA showing the interaction of Gβ subunit with K_V_7.2 or K_V_7.3 in HEK cells transfected with plasmids encoding for K_V_7.2 only, K_V_7.3 only, or K_V_7.2 + K_V_7.3, as indicated. The insets show a brightfield image of the cells. Nuclei (DAPI staining, blue) are also shown. Scale bar: 20 µm. (**B** and **C**) Scatter bar graphs showing the quantification of PLA puncta. Individual data points (n) represent one microscopic field; “n” were collected in 3-4 experimental sessions. (**B**) Puncta/cell for the K_V_7.2-Gβ pair (7.2 vs. 7.3, P < 0.0001; 7.3 vs. 7.2/7.3, P = 0.0005). (**C**) Puncta/cell for the K_V_7.3-Gβ pair (7.2 vs. 7.3, P = 0.0458; 7.2 vs. 7.2/3, P = 0.0011). Both subunits show colocalization with Gβ-containing Gβγ in intact cells.

We next investigated whether K_V_7–Gβ proximity was affected by the small-molecule Gβγ inhibitor gallein. We incubated HEK293 cells co-expressing K_V_7.2 and K_V_7.3 with 100 μM gallein for 10 or 30 min before PLA detection of K_V_7.2–Gβ or K_V_7.3–Gβ proximity (Fig. S5A). Gallein did not significantly alter the number of K_V_7.2–Gβ or K_V_7.3–Gβ PLA puncta compared to vehicle-treated cells. In contrast, under the same conditions, gallein decreased KCNQ4–Gβ PLA puncta by approximately 50%, consistent with previous findings (*16*) (Fig. S5B,C). We note that gallein binds to Gβ, occluding part of Gβ–effector interaction surfaces and selectively disrupting a subset of Gβγ–effector interactions. However, it spares some prominent Gβγ effectors such as GIRKs and Ca_V_2.2 (*47*). The differential effect of gallein on K_V_7.4–Gβ versus K_V_7.2/7.3–Gβ PLA signals is therefore consistent with distinct Gβγ-contact interfaces or distinct conformational arrangements of Gβγ relative to K_V_7.2/7.3 and K_V_7.4. Overall, these findings indicate that K_V_7.2 and K_V_7.3 reside in close proximity to Gβ in resting cells and that this proximity is retained in the presence of gallein.

### Peptide arrays identify multiple conserved intracellular K_V_7.2/7.3 regions that bind Gβγ

Having established that Gβγ lies in proximity to KV7.2/7.3 and regulates channel’s function and surface expression, we tested the hypothesis that Gβγ directly interacts with K_V_7.2 and K_V_7.3 subunits and mapped the regions of the channel that mediate the interaction with Gβγ. We generated peptide arrays of overlapping 25-mer peptides covering the cytosolic domains of K_V_7.2 and KV7.3, immobilized on nitrocellulose membranes. The membranes were incubated with purified Gβγ and probed with an anti–Gβ antibody to detect bound Gβγ (Fig. 5).

**Fig. 5.**
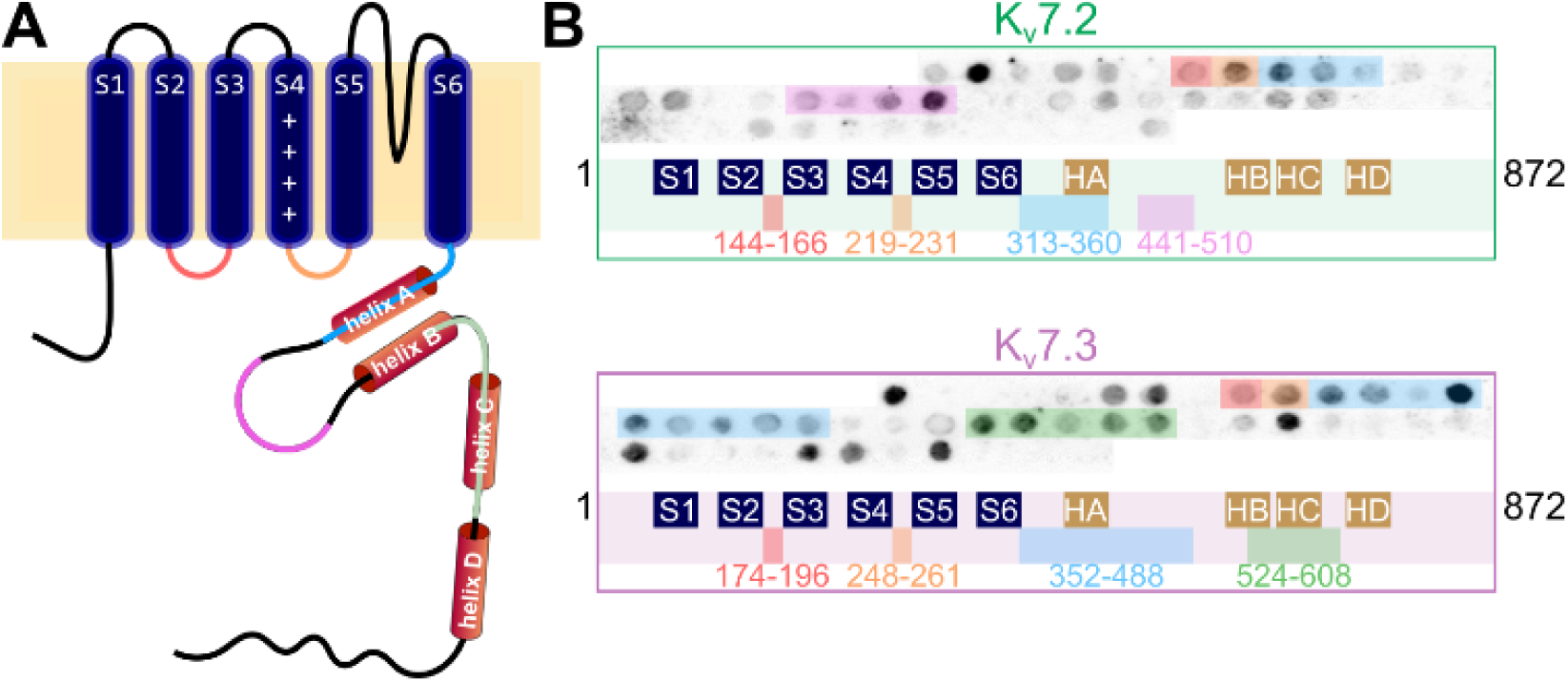
Peptide arrays analysis identifies multiple intracellular K_V_7.2/7.3 regions that bind Gβγ. (**A**) Schematic of a K_V_7 subunit (including S1-S6, S2-S3 and S4-S5 linkers, and C-terminal helices A-D) with intracellular regions color-coded according to the binding clusters in (B). (**B**) Peptide array membranes covering the complete intracellular sequences of human K_V_7.2 (top) and K_V_7.3 (bottom) probed with purified His-Gβγ. Darker spots indicate stronger binding. Bars and topology maps mark the principal clusters: K_V_7.2 residues 144-166 (S2-S3), 219-231 (S4-S5 linker), 313-360 (proximal C-terminus/helix A), and 441-510; K_V_7.3 residues 174-196, 248-261, 352-488, and 524-608 (helix B-C).

Rather than binding a single site, Gβγ bound multiple discrete regions distributed throughout the intracellular portions of both channel subunits (Fig. 5B; Fig. S6). For K_V_7.2, Gβγ– binding peptides mapped to residues 144–166 (S2-S3 linker), 219–231 (S4-S5 linker), 313–360 (proximal C terminus), and 441–510 (helices A-B region). For KV7.3, Gβγ bound to the homologous regions spanning residues 174–196, 248–261, 352–488, and 524–608. These Gβγ– binding regions were largely conserved between the two subunits, clustering within the S2-S3 linker, the S4-S5 linker, and the proximal C-terminal helices A-D (Fig. 5A, Fig. S6).

To place these results in a structural context, we generated an AlphaFold model of the K_V_7.2/7.3 heterotetramer (2:2 stoichiometry) in complex with Gβγ (Fig. S7). The model was colored according to per-residue confidence (pLDDT). The transmembrane and proximal C-terminal regions of K_V_7.2/7.3 were predicted with high to very high confidence, whereas parts of the N terminus and more distal C-terminal regions were predicted with lower confidence (Fig. S7A). Most regions identified as Gβγ–interacting in the peptide arrays localized to the Gβγ– contacting surface in the model (Fig. S7B-D). This finding supports the possibility that these conserved determinants contribute to the Gβγ-interaction interface of K_V_7.2/7.3.

### A disease–associated GNB1 mutation selectively impairs K_V_7.2/7.3 potentiation

Finally, we tested whether disease–causing mutations in *GNB1*, which encodes Gβ₁, affect M-current modulation. We examined the K78R, I80N, and I80T variants, all of which have previously been shown to alter Gβγ regulation of GIRK channels (*35, 37*). In our earlier work in GMPs, K78R exhibited WT-like surface expression at saturating Gβγ and increased expression at low Gβγ doses, together with a gain of function for GIRK2 channels. In contrast, I80N and I80T showed reduced surface expression and loss of function for GIRK2 channels (*37*). To obtain Gβ variant expression levels comparable to those of Gβ_WT_γ, we injected 5 ng Gβ_WT_ or Gβ_K78R_ RNA, 10 ng of Gβ_I80N_ and Gβ_I80T_ RNA, and Gγ RNA to maintain 5:1 Gβ:Gγ ratio.

Wild-type Gβγ and the K78R and I80T Gβ variants each significantly potentiated K_V_7.2/7.3 currents relative to channels expressed without Gβγ (Fig. 6A, B). In contrast, the I80N variant failed to enhance the current, which remained indistinguishable from the no-Gβγ condition and significantly different from the other Gβ variants (Fig. 6B). None of the variants altered the V_50_ (Fig. 6C) or K_a_ (Fig. 6D).

**Fig. 6.**
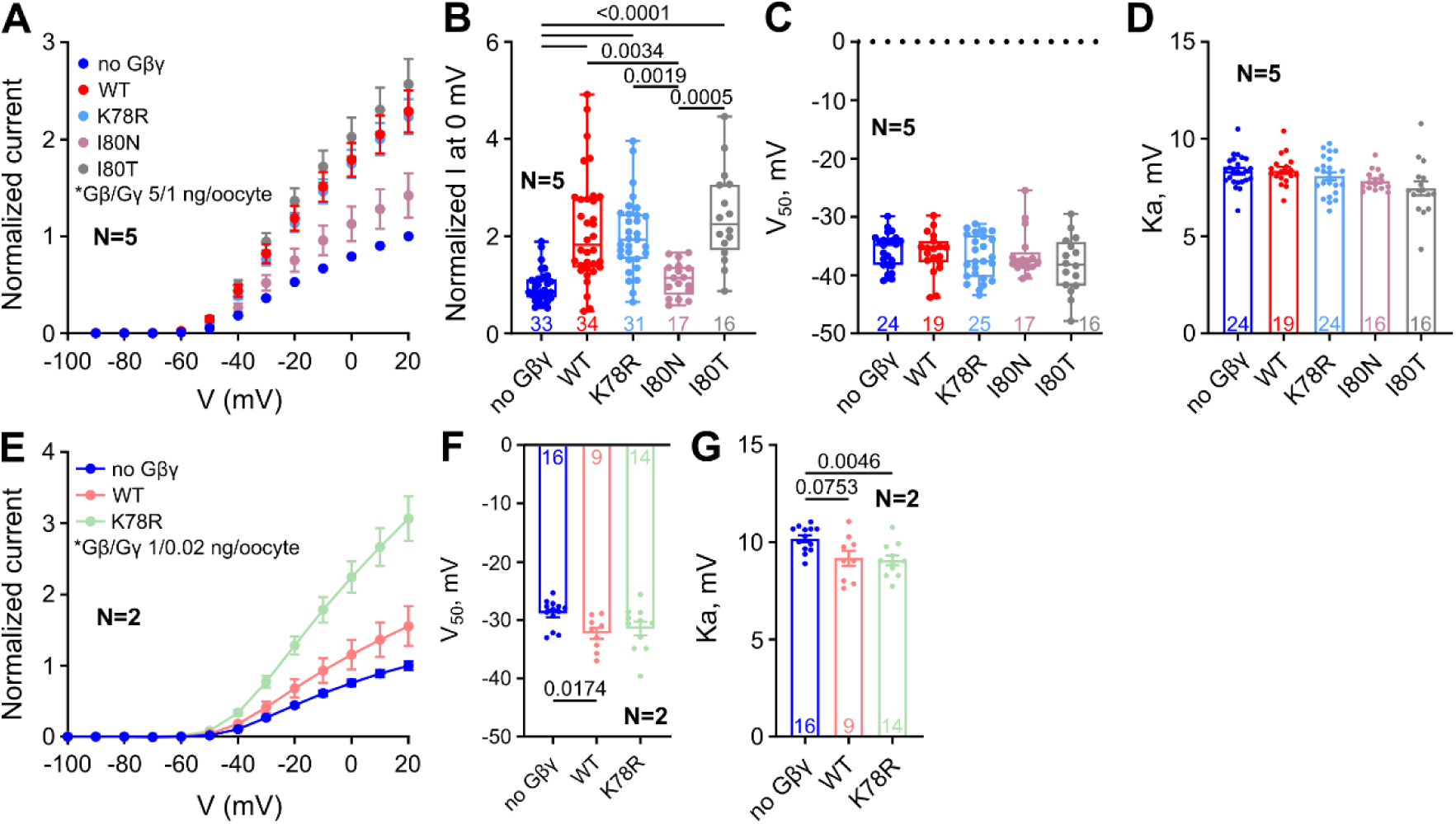
The GNB1 encephalopathy variants alter Gβγ-induced regulation of K_V_7.2/7.3. K_V_7.2/7.3 was expressed without Gβγ or with Gγ_2_ together with either WT Gβ_1_ or the variants K78R, I80N, or I80T. (**A**) Mean normalized I-V relationships from 5 experiments. Numbers of cells (n) are shown in B. (**B**) Normalized current at 0 mV: WT, K78R, and I80T activated K_V_7.2/7.3 current (WT, P < 0.0001; K78R, P = 0.0034; I80T, P = 0.0005), whereas I80N failed to enhance the current. (**C**, **D**). Gating parameters V_50_ (C) and K_a_ (D) were unchanged across variants. (**E**) Mean normalized I-V relationship with a lower dose of Gβγ (Gβ:Gγ RNA, 1:0.2 ng/oocyte). Comparison of normalized currents at 0 mV by one-way ANOVA showed that K78R activated K_V_7.2/7.3 current (p<0.0001), whereas, at this dose, the WT Gβγ does not significantly increase the M-current. (**F, G**) Gating parameters V_50_ and K_a_ show modest changes after expression of GβWTγ or Gβ_K78R_γ in this set of experiments.

Since Gβ_K78R_ showed gain-of-function behavior toward GIRK2 at low amounts of injected Gβγ RNAs (*35*), we examined its effect with K_V_7.2/7.3 under the same conditions (Gβ:Gγ, 1:0.2 ng/oocyte). At this dose, K78R potentiated K_V_7.2/7.3 currents more strongly than WT Gβγ (Fig. 6E) and produced a significant reduction in the slope factor K_a_ relative to the no-Gβγ condition; this effect was not significant for WT Gβγ (Fig. 6H). There was a modest hyperpolarizing shift in V_50_, which reached statistical significance for WT Gβγ, but not for Gβ_K78R_, in this set of experiments (Fig. 6F). The enhanced potentiation is consistent with K78R acting a gain-of-function mutant toward K_V_7.2/7.3, as previously found for GIRKs (*37*). Together, these results identify I80N as a selective loss-of-function mutation for KV7.2/7.3, reveal complementary gain-of-function behavior of K78R, and link Gβγ–dependent M-current regulation to *GNB1*– associated disease.

## Discussion

We identify Gβγ as a potent positive regulator of K_V_7.2/7.3 channels underlying the neuronal M-current and present convergent evidence for a direct channel–Gβγ interaction. Functional studies in *Xenopus* oocytes support a model in which Gβγ regulates the M-current at two levels: by increasing the abundance of the KV7.2/7.3 channels in the PM and by strengthening channel–PIP_2_ coupling. Furthermore, we present evidence that ambient Gβγ, presumably associated with the channel, is important for maintaining the basal activity of K_V_7.2/7.3. The GNB1 encephalopathy–associated I80N Gβ_1_ variant selectively abolishes M-current potentiation, whereas the K78R variant behaves as a gain-of-function, linking Gβγ-dependent regulation of K_V_7.2/7.3 to human neurodevelopmental disease.

The most striking effect of Gβγ on K_V_7.2/7.3 in our model system was an approximate doubling of whole-cell current and G_max_ upon Gβγ coexpression. Equally dramatic was the 60-80% decrease in basal current caused by Gβγ scavengers myr-phosducin and GRK2. Part of these bidirectional changes in M-current amplitude was likely driven by corresponding changes (increase and decrease, respectively) in PM abundance of channel subunits, as demonstrated by immunocytochemistry in giant excised membrane patches (GMPs). Changes in channel gating may also contribute, as discussed below. The coordinated increase in surface levels of the two subunits on addition of Gβγ (Fig. 2) suggests that overall channel stoichiometry remains unchanged at the population level. We note that this approach cannot resolve the effect of Gβγ on the subunit stoichiometry of individual heteromers – which remains debatable – with K_V_7.2: K_V_7.3 ratios ranging from 3:1 to 1:3 (*48–50*). Resolving this will require direct structural approaches, such as single-particle cryo-EM of K_V_7.2/7.3 in the presence and absence of Gβγ. Further, the GMP measurements quantify steady-state PM abundance and do not distinguish altered trafficking, retention, recycling, or degradation of channel subunits; the exact mechanisms underlying the Gβγ–induced enhancement of channel PM density remain to be elucidated.

Our results indicate that Gβγ regulation extends beyond channel surface abundance to functional channel–PIP_2_ coupling and, potentially, channel gating. The most evident gating effect was a slowed M-current decay upon PIP_2_ depletion via Ci-VSP, alongside the opposite effect observed upon Gβγ scavenging with GRK2 (Fig. 3). These findings strongly suggest that Gβγ stabilizes the interaction of PIP_2_ with the K_V_7.2/7.3 channel. Although an effect of Gβγ on Ci-VSP itself cannot be fully excluded, the reciprocal effects of Gβγ coexpression and Gβγ scavenging, together with their marked effects on Kᵥ7.2/7.3 current in the absence of Ci-VSP, support altered functional channel–PIP_2_ coupling as the most parsimonious interpretation. Because PIP_2_ is an obligatory cofactor for M-channel functioning and the principal target of G_q_-coupled receptor–mediated M-current suppression (*20, 51, 52*), a Gβγ–dependent increase in coupling to PIP2 has clear physiological implications, potentially rendering the M-current more resistant to PIP_2_ depletion downstream of G_q_/PLC signaling. A synergistic interplay between PIP_2_ and Gβγ was also reported for KV7.4 (*25*), raising the possibility that modulation by Gβγ of channel–PIP_2_ coupling is a conserved mechanism in the K_V_7 family, conceptually analogous to how Gβγ strengthens the PIP_2_ interaction with GIRK channels (*53, 54*).

In K_V_7 channels, improving or weakening coupling to PIP_2_ is expected to hyperpolarize or depolarize, respectively, the voltage dependence of channel opening (*55, 56*). At the first glance, in our experiments the voltage dependence of KV7.2/7.3 was not markedly affected by coexpression of Gβγ, with a small hyperpolarizing shift in V_50_ that reached statistical significance only in a portion of experiments (Figs. 1, 2, 6F). However, scavenging Gβγ with myr-phosducin drove V_50_ in the opposite direction. The overall shift in voltage dependence of K_V_7.2/7.3 between scavenger–expressing and Gβγ–expressing cells was approximately 6 mV (Fig. 2M), comparable to the 6-10 mV shift produced by Gβγ in K_V_7.4 (*16, 57*). Together, these results indicate that Gβγ not only regulates the channel’s PM abundance but also gating, rendering KV7.2/7.3 more sensitive to depolarization. Additionally, PIP_2_ is known to increase the maximal open probability (P_o,max_) of M-channels, even at saturating voltages (*58*). Consequently, Gβγ– induced alteration in PIP_2_ coupling and the resulting changes in the voltage dependence and Po,max are expected to contribute to the observed changes in M-current amplitude upon Gβγ expression or scavenging. The exact relative contribution of changes in gating compared to changes in PM abundance is unclear and awaits further study.

A further important aspect of Gβγ regulation is highlighted by the observed dramatic decrease in M-current caused by coexpression of Gβγ scavengers, myr-phosducin and GRK2. This finding suggests that Gβγ is required to maintain basal channel activity. An analogous dependence of basal activity on ambient Gβγ was reported for K_V_7.4, and it has been proposed that association of Gβγ with K_V_7.4 is necessary for proper response to voltage (*59*). A similar mechanism has been shown to maintain the basal activity of the GIRK1/2 channels (*10*). Such “basal” association does not need to involve a pre-existing non-dissociable complex; rather, it is dynamic, involving reversible protein–protein interactions (*39*). PLA further supports nanoscale proximity between Gβγ and K_V_7.2/7.3 in intact cells, although it cannot, by itself, distinguish direct binding from close association within a larger protein complex. With this reservation, the PLA results are compatible with Gβγ and K_V_7.2/7.3 association in resting state and without any GPCR activation. In summary, our current and previous findings extent the importance of channel-associated Gβγ for proper voltage-dependent gating from K_V_7.4 to KV7.2/7.3.

Importantly, non-prenylated Gβγ_C68S_ failed to regulate KV7.2/7.3 current and PM abundance, establishing that Gβγ action requires its membrane attachment and is, therefore, membrane–delimited. Such regulation, in which prenylated, membrane–anchored Gβγ directly engages the cytosolic regions of an integral membrane protein, is a hallmark of direct regulation of classic Gβγ targets such as GIRK and Ca_V_2 channels and some adenylyl cyclases (*9, 60*). A direct interaction of Gβγ with both K_V_7.2 and K_V_7.3 subunits is supported by the peptide array assay, in which purified Gβγ bound 25-mer peptides representing channel segments immobilized on a solid support.

The peptide arrays also allowed us to map the candidate Gβγ–binding regions in K_V_7.2 and K_V_7.3. In parallel, we used an alternative approach, with a computational model constructed using AlphaFold. Both the peptide arrays and the AlphaFold model identified similar Gβγ–interacting segments: parts of the C-terminus and the S2-S3 and S4-S5 linkers, at or near the binding sites of other important regulators. Thus, calmodulin binds at or near several of the mapped Gβγ-contact segments in the C-terminal helices (*23*). The S2-S3 and S4-S5 linkers are known to anchor and functionally interact with PIP_2_ (*22, 55*), raising the possibility that this could be part of the mechanism by which Gβγ allosterically reinforces PIP_2_–channel interactions.

Our screening of pathogenic GNB1 mutant variants links this regulatory phenomenon to human disease. The I80N variant showed loss-of-function, failing to potentiate K_V_7.2/7.3, whereas K78R retained, and at low doses exceeded, WT potentiation, acting essentially as a gain-of-function mutant (Fig. 6). Importantly, loss-of-function and, more rarely, gain-of-function KCNQ2 mutations are strongly associated with developmental epileptic encephalopathies (*61–63*). Correspondingly, loss-or gain-of-function mutations in an M-channel stimulator – in this case Gβγ – could be expected to have to have analogous effects on neuronal excitability. Interestingly, the same mutations have been shown to have a similar effect on Gβγ-dependent activation of GIRK channels (*35, 37*), another K^+^ channel that, like the M-current, dampens neuronal excitability. In summary, our findings identify altered M-current regulation as a potential contributor to GNB1 encephalopathy, alongside previously described GIRK dysregulation, and support evaluation of KCNQ-targeting pharmacotherapies for GNB1 encephalopathy-related epilepsy.

### Limitations of the study

The peptide arrays used here to map the potential Gβγ-interacting channel segments detect linear peptide determinants and are prone to false positives at basic or charged stretches; the mapped segments should therefore be treated as candidate contact sites pending orthogonal validation by solution–phase binding. Structure–guided point mutations that abolish potentiation may seem like a promising direction. However, in the conserved regions that we have identified, any change may impair channel function or its interaction with other partners, particularly PIP_2_ and calmodulin. We note that the peptide array results were corroborated by the AlphaFold analysis. Although the latter remains a computational model and requires structural confirmation, the similarity between the Gβγ–binding segments identified by the two independent methods lends credibility to these results.

Our study used *Xenopus* oocytes as a heterologous expression system for all functional tests (only PLA was conducted in HEK cells, which are of human origin). Trafficking and degradation of expressed proteins may be regulated differently than in mammalian cells and, specifically, neurons, urging for caution in generalizing the effect of Gβγ on PM abundance of K_V_7.2/7.3. On the other hand, oocytes usually faithfully reproduce the gating properties of ion channel and their regulation by interactors and remain among the most widely used and best-characterized model systems for gating studies. Looking ahead, testing the effect of Gβγ on K_V_7.2/7.3 channels in neurons would be highly informative for establishing the physiological relevance of this interaction.

## Methods

### Ethical approval of Xenopus laevis, oocyte preparation, and electrophysiology

Experiments were approved by Tel Aviv University Institutional Animal Care and Use Committee (permit #01-20-083). Maintenance and surgery of female frogs were as described previously (*64*). In brief, adult female *Xenopus laevis* frogs were purchased from Xenopus 1 (Dexter, MI). For surgery, frogs were anesthetized in 0.2% tricaine methanesulfonate (MS-222). After collecting oocytes from the ovary, frogs were sutured and allowed to recover from anesthesia. Then, frogs were returned to a separate tank for postoperative animals. Oocytes were defolliculated in Ca^2+^-free solution (in mM: 96 NaCl, 2 KCl, 1 MgCl_2_, 5 HEPES, pH 7.5) with 1-2 mg/mL collagenase (Sigma, Type 1A). Two hours later, oocytes were washed with NDE solution (in mM: 96 NaCl, 2 KCl, 1 MgCl2, 1 CaCl_2_, 5 HEPES, 2.5 mM sodium pyruvate, 50 µg/ml gentamycin, pH 7.5) (*64*) and let to rest for at least 2 hours. Then, oocytes were injected with RNA and incubated for 3 days at 20–22◦C in NDE solution.

### DNA constructs and cRNA preparation

Coding sequences of all DNA constructs used for protein expression in Xenopus *oocytes* (Table S2) were inserted into pGEM-HE, pGEM-HJ or pXOOM vectors that contain 5’ and 3’ untranslated regions from *Xenopus* β-globin (*65*). GNB1 (Gβ_1_), GNG2 (Gγ_2_), myristoylated phosducin (myr-phosducin), the prenylation-deficient Gγ2 mutant (Gγ2-C68S), the disease-associated Gβ_1_ variants K78R, I80N, and I80T, and GIRK subunits 1 and 2 (GIRK1 and GIRK2) were described in previous publications (*39, 43, 66*). Human KCNQ2 (K_V_7.2) and KCNQ3 (K_V_7.3) in pGEM-HJ plasmid were kind gifts from Rene Barro-Soria (University of Miami, USA). N-terminally tagged human YFP-KCNQ3, human KCNQ4 (K_V_7.4) were in pXOOM vector. Full-length G-protein kinase 2 (GRK2) in pXOOM was a kind gift from Kristoffer Sahlholm (Umeå University, Sweden). N-terminal HA-and Myc-tags were introduced at positions that do not interfere with channel assembly or function of KCNQ2 and KCNQ3, using short linkers (HA-KCNQ2 and Myc-KCNQ3) (Table S2). *Ciona intestinalis* voltage-sensing phosphatase (Ci-VSP) was purchased from Addgene (Watertown, MA, USA) and subcloned into pGEM-HJ using LigON kit (E1050-01, EURx Molecular Biology Products). mRNAs were prepared as described previously (*67*). Capped cRNA was synthesized in vitro with T7 RNA polymerase from linearized plasmid, purified, and quantified spectrophotometrically; integrity was confirmed by agarose gel electrophoresis. Gβ and Gγ RNAs were injected at 5 and 1 ng per oocyte, respectively, which were empirically optimized for functional expression and robustly activated GIRK channels expressed in the oocytes (*39*). Amounts of Kᵥ7.2 and Kᵥ7.3 RNAs ranged from 0.5 to 2 ng each, and other RNAs have been varied to produce optimal effects and are indicated in text and figure legends.

### Two-electrode voltage clamp (TEVC) in oocytes: electrophysiology and analysis

For electrophysiological experiments, we used two-electrode voltage clamp (TEVC). K_V_7 channel currents were recorded in ND96 solution (low K^+^) (in mM: 96 NaCl, 2 KCl, 1 MgCl_2_, 1 CaCl_2_, 5 HEPES, pH 7.5) at 20–22◦C. GIRK currents were measured in high-K^+^ (HK24) solution (in mM: 24 KCl, 74 NaCl, 1 MgCl2, 1 CaCl2, 5 HEPES, pH adjusted to 7.5 with KOH; 30 s) (*64*). Whole-cell currents were measured using a GeneClamp 500B amplifier and an Axon Digidata 1440a (Molecular Devices). Holding potential was -90 mV. Currents were acquired at 200-2,000 Hz and filtered at 20 Hz. Recording was initiated upon stabilization of the resting membrane potential. Experiments were performed using the following voltage protocols (→ represents proceeding to the next epoch; see Fig. S8):

1. Voltage step protocol sequence (for I-V relationships): -90 mV (200 ms) → -120 mV (200 ms) → -90 mV (50 ms) → -100 mV to +20 mV in 10 mV increments (3000 ms) → -60 mV (3000 ms) → -90 mV (200 ms). (Fig. S8A).
2. To deplete PIP_2_ by activation of Ci-VSP, we used the following PIP_2_ depletion protocol: -90 mV (200 ms) → -120 mV (200 ms) → -90 mV (50 ms) → +20 mV (15000 ms) → -60 mV (6000 ms) → -90 mV (200 ms). (Fig. S8B).
3. For the recording of GIRK currents the voltage was held at -80 mV throughout the whole recording, and the indicated solutions were sequentially applied: ND96 (5 s) → Voltage ramp from -120 to 50 mV (2 s) → HK24 solution → Voltage ramp from - 120 mV to 50 mV (2 s) → HK24 + GIRK blocker (2.5 mM Ba^2+^) (Fig. S8C).

Current amplitudes were measured at the end of the test pulse per each voltage for the I-V curve, and at 0 mV for cross-condition comparison. Leak was calculated using a linear fit extrapolated from the part of the I-V relationship where M-channels were deactivated (−100 mV : -70 mV), then the leak was subtracted from the test pulse per each voltage of the I-V curve. Maximal conductance (G_max_) was derived from fitting of the I-V relationship to Boltzmann I-V equation, where the K^+^ current reversal potential (V_rev_) was assumed to be -95 mV:

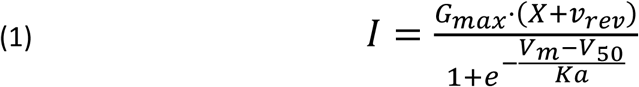

Conductance-voltage (G-V) relationships were constructed from tail-current amplitudes measured 45 ms after the start of repolarization at a fixed repolarizing potential (−60 mV), rather than from end-of-pulse currents, so that the driving force was constant and V_rev_-independent across test voltages. G/G_max_ value was obtained by normalizing I_tail_ to the largest Itail in the I-V relationship, and the G-V relationship was fitted to Boltzmann equation in the form:

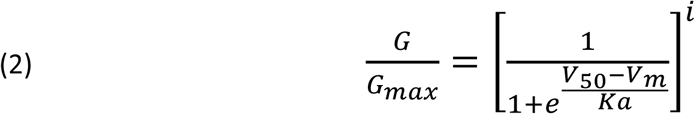

and, where indicated, raised to the first (*i* = 1) or fourth (*i* = 4) power. The fourth-power assumption follows the classic Hodgkin–Huxley *n*⁴ formalism (*41*), in which channel opening requires the independent and identical activation of four gating particles such that the pore conducts only after all four sensors have activated (*68, 69*). Fitting yielded the half-activation voltage (*V*_50_) and slope factor (*K*_a_). Deactivation kinetics were quantified by fitting tail-current decay with a double-exponential function to obtain fast (τ_1_) and slow (τ_2_) time constants and their relative amplitudes (A_2_/A_1_):

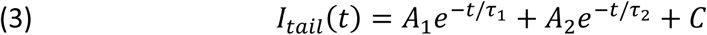

### Giant membrane patches (GMPs)

GMPs were prepared and imaged as described (*39*). Oocytes were devitellinized using tweezers in hypertonic solution (in mM: 6 NaCl, 150 KCl, 4 MgCl2, 10 Hepes, pH 7.6). The devitellinized oocytes were transferred onto a Thermanox^TM^ coverslip (Nunc, Roskilde, Denmark) and immersed in Ca^2+^-free ND96 solution with their black hemisphere facing the coverslip, for 30–45 min. The oocytes were then suctioned using a Pasteur pipette, leaving a GMP attached to the coverslip, with the cytosolic part facing the medium. The coverslip was washed thoroughly with fresh ND96 solution and fixated with 4% formaldehyde for 30 min. Fixed GMPs were blocked in 5% milk in in Tris-Buffered Saline with Tween 20 (TBST), and nonspecific binding was further blocked with donkey IgG 1:200 (Jackson ImmunoResearch, West Grove, PA, USA). Primary rabbit anti-Gβ (1:500; GeneTex, GTX114442), rabbit anti-GFP (1:500; Abcam, ab6556), HA-probe Antibody (F-7) Alexa Fluor® 647 conjugated (1:200; Santa Cruz, sc-7392 AF647), Myc Antibody (9E10) Alexa Fluor® 488 conjugated (1:200; Santa Cruz, sc-40 AF488) were applied for 1 hour with gentle shaking at room temperature. Then, for non-conjugated primary antibodies we applied Cy3 donkey anti-rabbit secondary antibody (1:400; Jackson, 711-165-152) for 30 min at room temperature, washed with TBST and mounted on a slide for visualization. Immunostained slides were kept at 4 °C in the dark.

### Confocal imaging in oocytes

Confocal imaging and analysis were performed as described (*37, 39*) with a Zeiss 710 META confocal microscope and BC43 - Benchtop Spinning Disc Confocal Microscope (Oxford Instruments Andor, Belfast, UK) in the Research Infrastructure Core Facilities (RICF) at the Faculty of Medical & Health Sciences in Tel Aviv University. Imaging of proteins in GMPs was performed in λ-mode as described (*37, 39*). Cy3 was excited using 543 nm HeNe laser and emission was collected at 556-615 nm. When proteins in GMPs were imaged using the BC43, Alexa Fluor 488, Cy3, and Alexa Flour 647 were excited with green (488 nm excitation, 517–541 nm emission), red (561 nm excitation, 580–610 nm emission), and far-red (638 nm excitation, 671–745 nm emission) lasers, respectively. For both Zeiss 710 and BC43, images centered on edges of the membrane patches, so that background fluorescence from coverslip could be seen and subtracted. Two regions of interest were chosen: one comprising most of the membrane patch within the field of view, and another comprising background fluorescence, which was subtracted from the signal obtained from the patch. The signal from GMPs prepared from native oocytes, immunostained using the same protocol, was subtracted from all groups.

### HEK293 cell culture and transfection

HEK293 cells were maintained in Dulbecco’s modified Eagle’s medium (DMEM) supplemented with 10% (vol/vol) FBS, 1% L-glutamine, 1% penicillin/streptomycin in a 37 °C incubator with 5% (vol/vol) CO2. For transfection, cells were plated on 12-mm diameter coverslips placed in 24-well plates; the day after plating, cells were transfected with different combinations of pcDNA3.1-KCNQ2, pcDNA3.1-KCNQ3, and pcDNA3.1-KCNQ4 (0.8 µg DNA per well), using lipofectamine 2000 (Life Technologies, Milan, Italy), according to manufacturer’s instruction. For pharmacological treatments, 24 h after transfection, cells were incubated for 10 or 30 min at 37°C with 100 µM gallein or vehicle (0.1% DMSO), both dissolved in DMEM.

### Proximity Ligation Assay

Cells were fixed with 4% (vol/vol) paraformaldehyde for 15 min at room temperature before permeabilization in 0.1% Triton X-100 for 5 min. PLA was performed using a DuoLink kit (Merck, Milan, Italy), according to manufacturer’s instructions, as previously described (*16*). Cells were blocked for 1 h at 37 °C in DuoLink blocking solution and incubated overnight at 4°C with mouse anti-Gβ (1:100 Santa Cruz Biotechnology, Heidelberg, Germany), and one of the following antibodies: rabbit anti-KV7.2 (1:100, GeneTex, Freising, Germany), rabbit anti-K_V_7.3 or anti-K_V_7.4 (both 1:100, Alomone Lab, Jerusalem, Israel). The following day, cells were incubated with DuoLink anti-rabbit PLUS and anti-mouse MINUS probes for 1 h at 37 °C. The hybridized oligonucleotides were ligated for 30 min at 37 °C before the rolling circle amplification, performed for 100 min at 37 °C. Coverslips were mounted using mounting medium with DAPI (Merck, Milan, Italy), and imaged using a confocal microscope (Zeiss 710 Meta). Images (7 per experimental field) were acquired over the Z-axis (Z-stack) using the Zeiss 710 Meta confocal microscope in 3-4 experimental sessions. The number of puncta was measured using FIJI software; number of puncta in each image acquired along the Z-axis was summed up and then divided by the number of nuclei in the experimental field, to obtain the “puncta/cell” value. Data were expressed as mean ± S.E.M. (standard error of the mean). Statistical analysis (one-way ANOVA with Tukey’s multiple comparison test) was performed using Prism 9.0 software.

### Gβγ expression and purification

We used His_6_-Gβγ and His_6_-Gβγ_C68S_ from the same preparation as that described in a previous publication (*39*). Gβ_1_ and Gγ_2_ were expressed in *Trichoplusia ni* (*T.ni*) cells. The His_6_-Gβ_WT_γ was extracted from the membrane fraction, which yielded a final purified preparation that was >95% prenylated (*70*). Protein purity was analyzed using SDS-PAGE and by Western blot using anti-Gβ_1_ and anti-His tag antibodies.

### Peptide spot array

Peptide arrays were generated by automatic SPOT synthesis and blotted on a Whatman membrane (*71*). Cytosolic segments of K_V_7.2 and K_V_7.3 were spot-synthesized as overlapping 25-mer peptides, shifted by 10 amino acids along the sequence, using AutoSpot Robot ASS 222 (Intavis Bioanalytical Instruments, Cologne, Germany). The peptides were designed according to human KCNQ2 (Uniprot: O43526) and human KCNQ3 (Uniprot: O43525). The interaction with spot-synthesized peptides was investigated by an overlay assay. Following blocking for 1 hour at room temperature with 5% BSA in 20 mM Tris and 150 mM NaCl with 0.1% Tween-20 (TBST), 0.016–0.16 μM purified His-Gβγ was incubated with the immobilized peptide-dots, overnight at 4 °C. His-Gβγ was detected by anti-GNB1 antibody (GTX114442) at 1:500 or 1:1000 dilution, followed by anti-rabbit HRP-coupled secondary antibody (1:40000) incubated with 5% BSA/TBST, and the membrane was imaged using Fusion FX7 (Vilber Lourmat, Collégien, France).

### Data analysis and statistics

Electrophysiological recordings were analyzed using pCLAMP (Molecular Devices) and custom software developed for this study and deposited in Zenodo (doi:10.5281/zenodo.22501858), using Python 3.10 and the pyABF package (pyABF). Statistical analyses were performed using GraphPad Prism 11 (GraphPad Software, San Diego, CA, USA). Outliers were identified using the ROUT method (Q=0.01Q=0.01) and excluded. Data are presented as mean ± SEM for normally distributed data and otherwise as box plots showing the median and minimum–maximum whiskers. For data that passed the Shapiro–Wilk normality test, unpaired t tests or one-way ANOVA with Tukey’s post hoc test were used. Otherwise, Mann–Whitney tests or Kruskal–Wallis tests with Dunn’s post hoc test were used.

## Acknowledgments

Funding:

This work was supported by grants from the Israel Science Foundation (ISF # 2780/20 to J.A.H., ISF #581/22 to N.D. and ISF # 1623/21 to I.L.), Vienna Science and Technology Fund (WWTF #LS25-014 to ND), British Heart Foundation (PG/25/12329 to I.G.), Deutsche Forschungsgemeinschaft (DFG; KL1415/13-1 and KL1415/14-1 to E.K.), Italian Ministry for University and Research (MUR PNR 2021-2027, Project MNESYS FORWARD to M.T.).

## Authors contributions

Conceptualization: N.D., B.S., I.G., I.L.

Methodology: N.D., B.S., V.B., J.A.H., E.K., I.G., I.L.

Investigation: B.S., I.P.-D., T.K-R., S.W., F.D.M., A.G.-S., K.Z.

Formal analysis: B.S., I.P.-D., T.K-R., S.W., F.D.M., V.B., N.D.

Resources: J.A.H., E.K., V.B., N.D.

Data curation: B.S.

Writing – original draft: B.S., V.B., N.D.

Writing – review & editing: B.S., V.B., I.G., I.L. N.D.

Supervision: J.A.H., M.T., V.B., I.G., I.L., N.D.

Funding Acquisition: J.A.H., M.T., V.B., E.K., I.G., I.L., N.D.

## Competing interests

M.T. and V.B. are partners of Keyon Lab, a startup company and University spinoff active in the area of drug development for neuropsychiatric diseases. All other authors declare no conflict of interest.

## Data, materials, and code availability

All reported data are presented in Results and Supplementary Materials. Original recordings and summary files will be made available upon reasonable request. Analysis routines are freely available on Zenodo (doi:10.5281/zenodo.22501858).

## Supplementary material for

### Contents

Supplementary Figures S1-S8 Supplementary Tables S1-S3

### Supplementary Figures

**Fig. S1.**
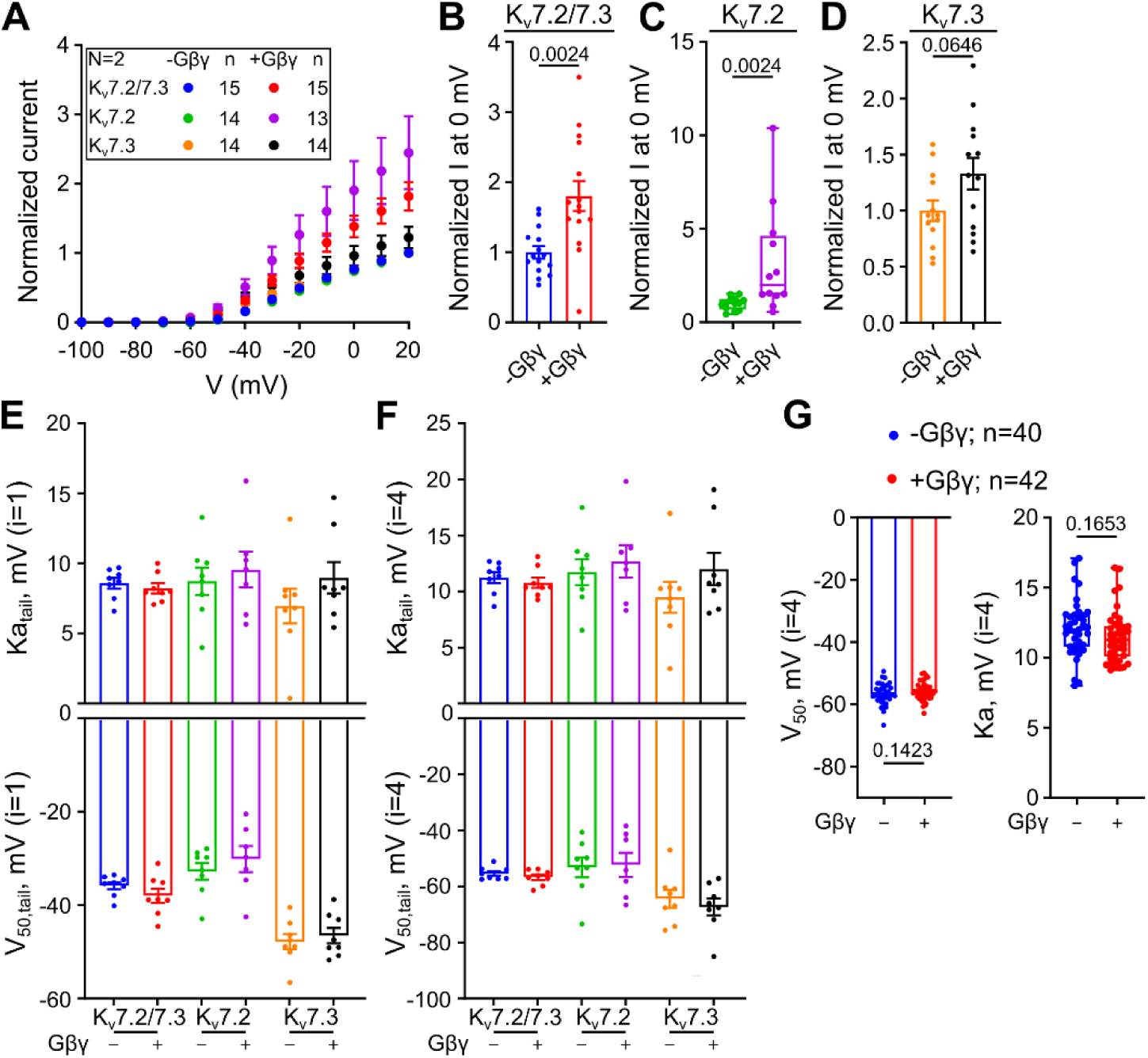
Gβγ potentiation of homomeric and heteromeric K_V_7 channels. (**A**) Mean I-V relationships for K_V_7.2/7.3, K_V_7.2, and K_V_7.3 with Gβγ, normalized to the average current at +20 mV in the control group of the same experiment (K_V_7.2/7.3, KV7.2, and K_V_7.3 without Gβγ, respectively). (**B**-**D**) Normalized current at 0 mV for K_V_7.2/7.3 (**B**, P = 0.0024), K_V_7.2 (**C**, P = 0.0024), and K_V_7.3 (**D**, P = 0.0646). (**E** and **F**) K_a_ and V_50_ obtained from tail current analysis in this set of experiments, for i = 1 (**E**) and i = 4 (**F**). (**G**), K_a_ and V_50_ obtained from tail current analysis in the set of experiments shown in Fig. 1, for i = 4.

**Fig. S2.**
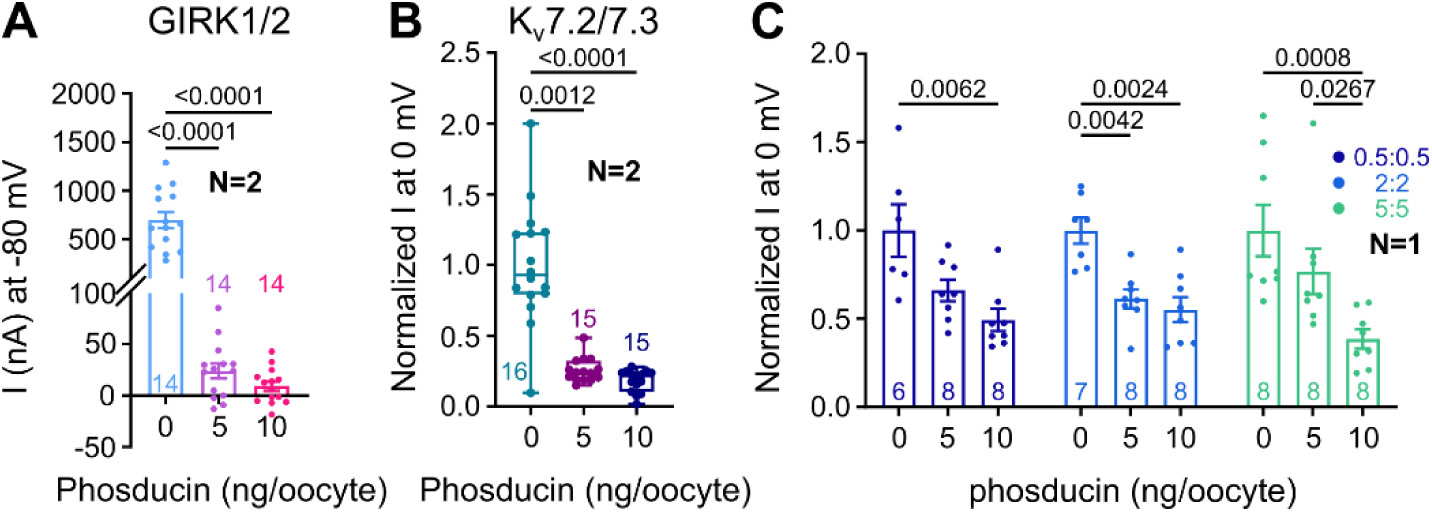
Dose-dependent suppression of channel function by the Gβγ scavenger myr-phosducin. (**A**) Myr-phosducin suppressed GIRK1/2 current measured at -80 mV (P < 0.0001 with 5 and 10 ng/oocyte of myr-phosducin RNA; RNA doses of GIRK1:GIRK2 were 0.05:0.05 ng/oocyte). (**B**) Myr-phosducin dose-dependently reduced basal K_V_7.2/7.3 current at 0 mV (5 ng, P = 0.0012; 10 ng, P < 0.0001; RNA doses of K_V_7.2:K_V_7.3 were 0.5:0.5 ng/oocyte). (**C**) The reduction of currents by myr-phosducin tales place in a range of K_V_7.2:KV7.3 RNA ratios (0.5:0.5, 2:2, 5:5 ng/oocyte).

**Fig. S3.**
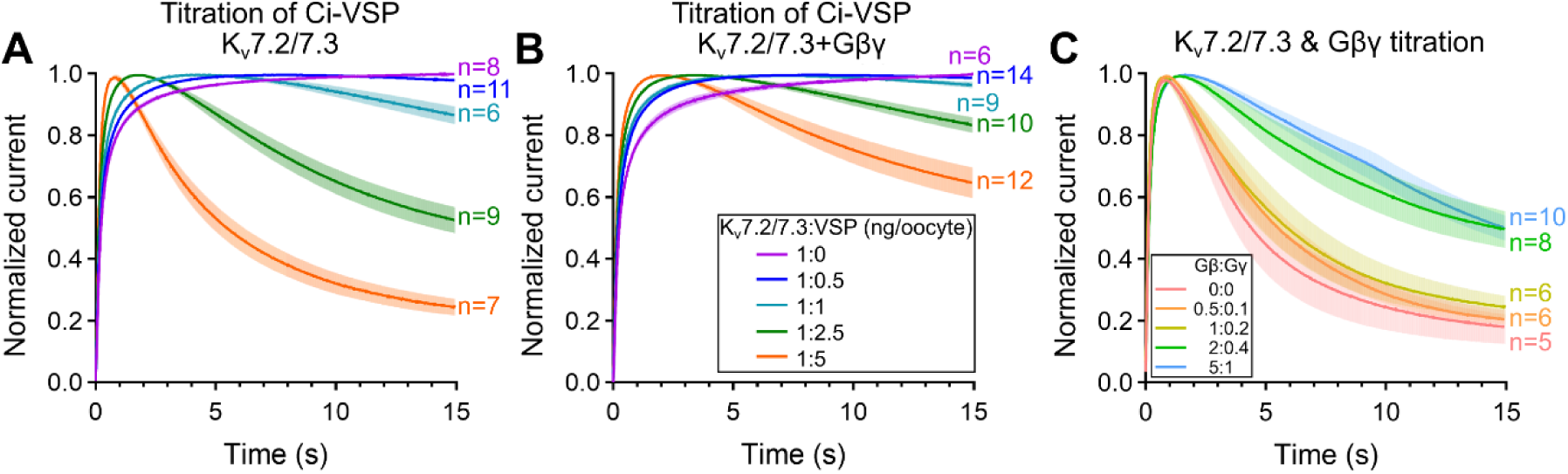
Ci-VSP/Gβγ titration of PIP2-depletion kinetics. (**A** and **B**) Averaged normalized current records at +20 mV, shown as mean±SEM, with varied K_V_7.2/7.3:Ci-VSP RNA ratios without (**A**) and with Gβγ (**B**). K_V_7.2/7.3 RNA dose was 1 ng of each subunit. (**C**) Increasing doses of Gβγ RNA progressive slow down the current decay. K_V_7.2/7.3 RNA dose was 0.5 ng/oocyte of each subunit.

**Fig. S4.**
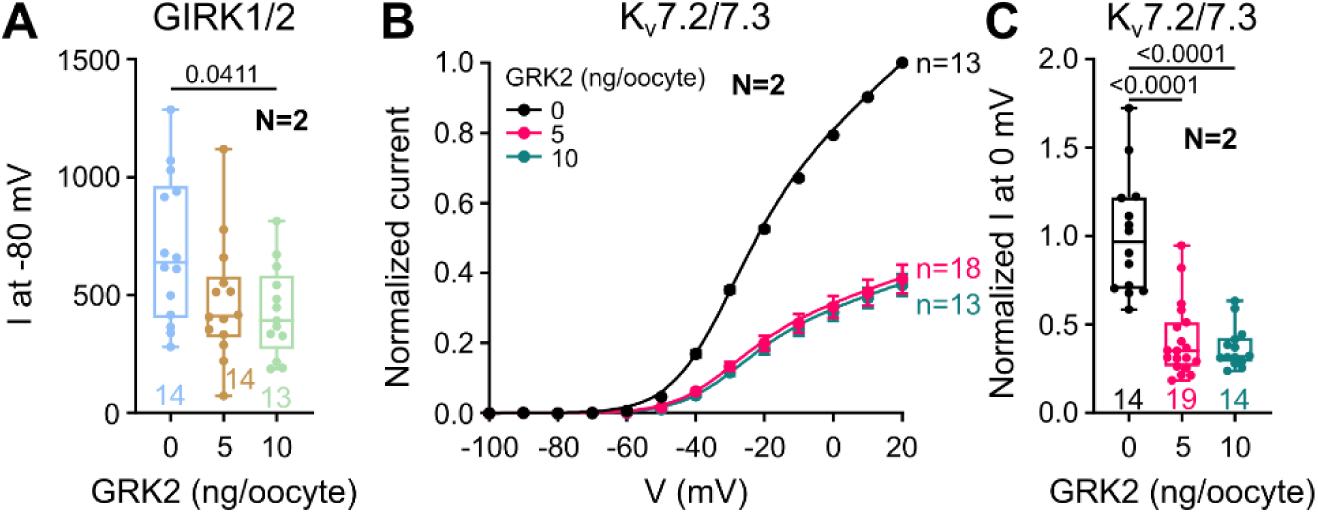
GRK2 acts as a Gβγ scavenger and reduces basal GIRK1/2 and K_V_7.2/7.3 current. (**A**) basal GIRK1/2 current measured at -80 mV as a positive control for Gβγ scavenging in high-K^+^ (24 mM) external solution. GRK2 was expressed at 5 and 10 ng/oocyte. Significant reduction was observed with 10 ng/oocyte RNA of GRK2 (P = 0.0411). (**B**) Mean normalized I-V relationships for K_V_7.2/7.3 with GRK2 at 0, 5, and 10 ng/oocyte. (**C**) Normalized K_V_7.2/7.3 current at 0 mV without or with GRK2 (5 or 10 ng RNA/oocyte). GRK2 reduced basal current at both RNA doses. Together these controls confirm that co-expressed GRK2 effectively sequesters Gβγ in oocytes, validating its use as a scavenger in this study.

**Fig. S5.**
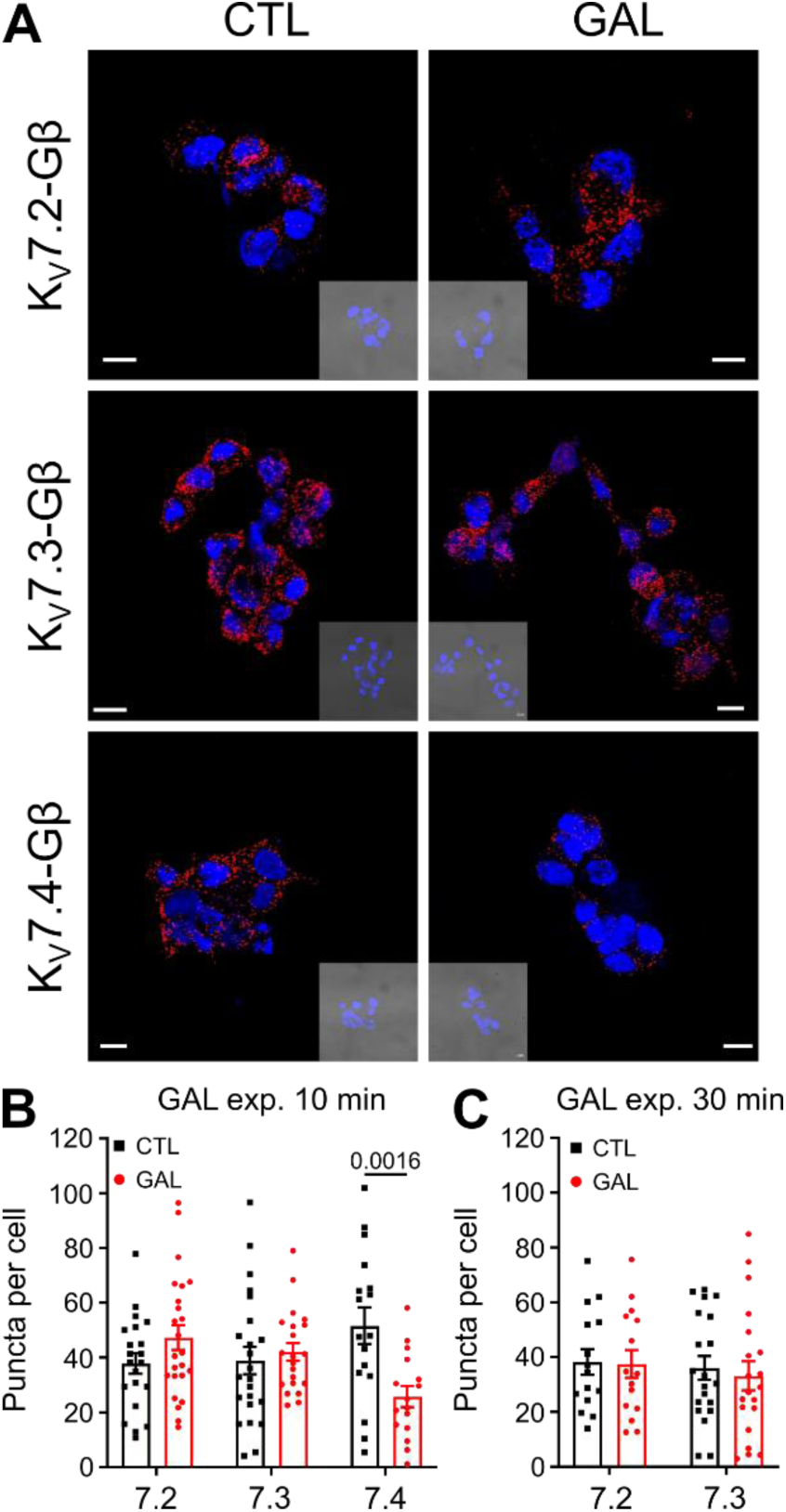
Gallein sensitivity confirms specificity of the channel-Gβ PLA signal. (**A**) Representative images of Proximity Ligation Assay (PLA) showing the interaction of Gβ subunit with K_V_7.2, K_V_7.3 or K_V_7.4 in HEK cells transfected with plasmids encoding for K_V_7.2 and K_V_7.3, or K_V_7.4, as indicated. The insets show a brightfield image of the cells. Nuclei (DAPI staining, blue) are also shown. Scale bar: 10 µm. (**B** and **C**) Scatter bar graphs shows the quantification of PLA puncta.Individual data points (n) represent one microscopic field; “n” were collected in 3-4 experimental sessions. **(B)** After 10-min gallein, the K_V_7.4-Gβ signal was reduced (*P* = 0.0016) without affecting K_V_7.2 or K_V_7.3. **(C)** A 30-min exposure produced no significant change for K_V_7.2 or K_V_7.3.

**Fig. S6.**
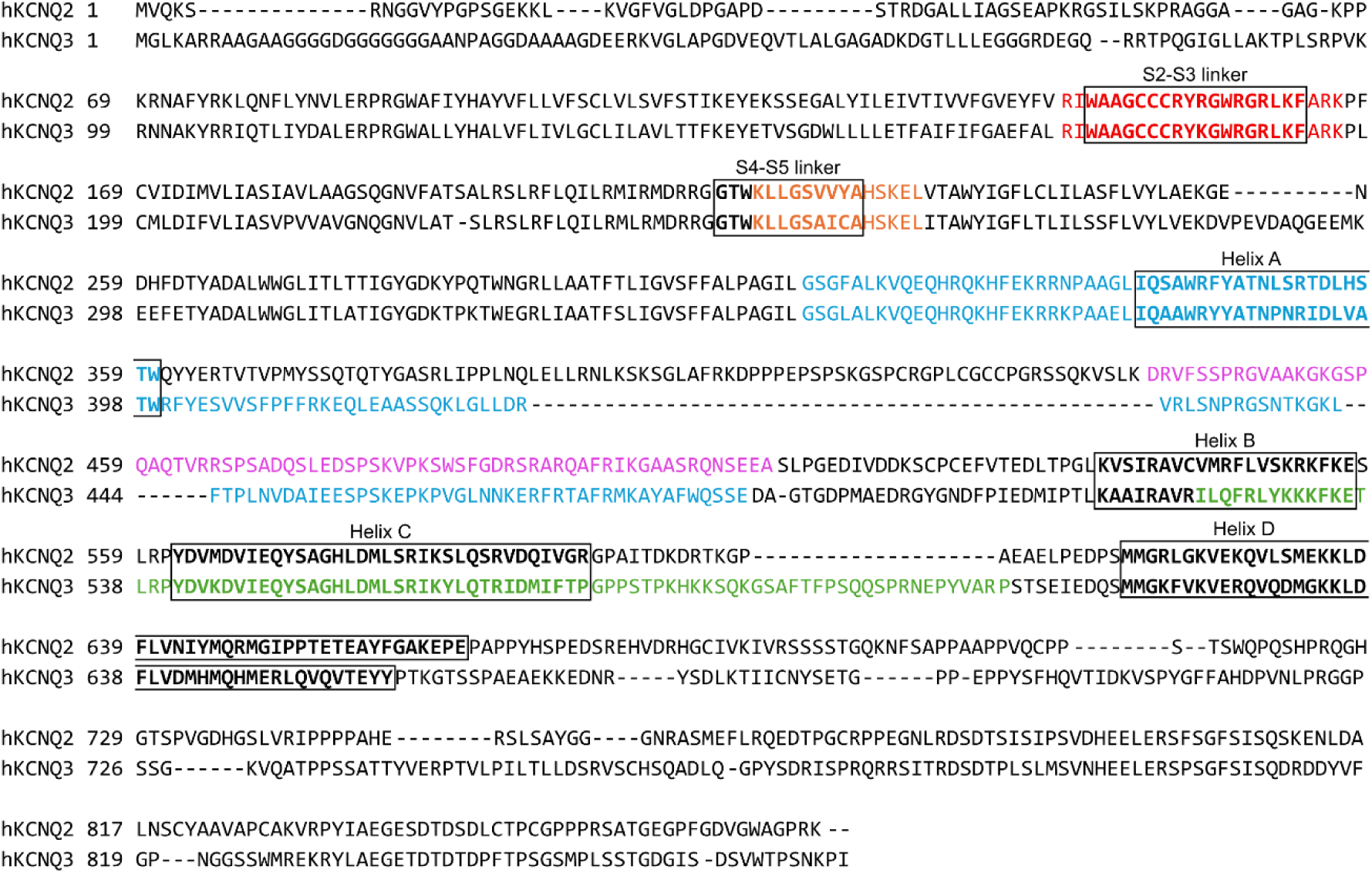
Sequence conservation of the Gβγ-binding regions between human K_V_7.2 and K_V_7.3. Pairwise alignment of human K_V_7.2 and K_V_7.3 intracellular sequences; the S2-S3 and S4-S5 linkers and C-terminal helices A-D are annotated with black rectangles and the binding clusters identified by peptide array analysis from Fig. 5 are highlighted in the same color code as in Fig. 5. The principal determinants are located in conserved or homologous regions of the two subunits.

**Fig. S7.**
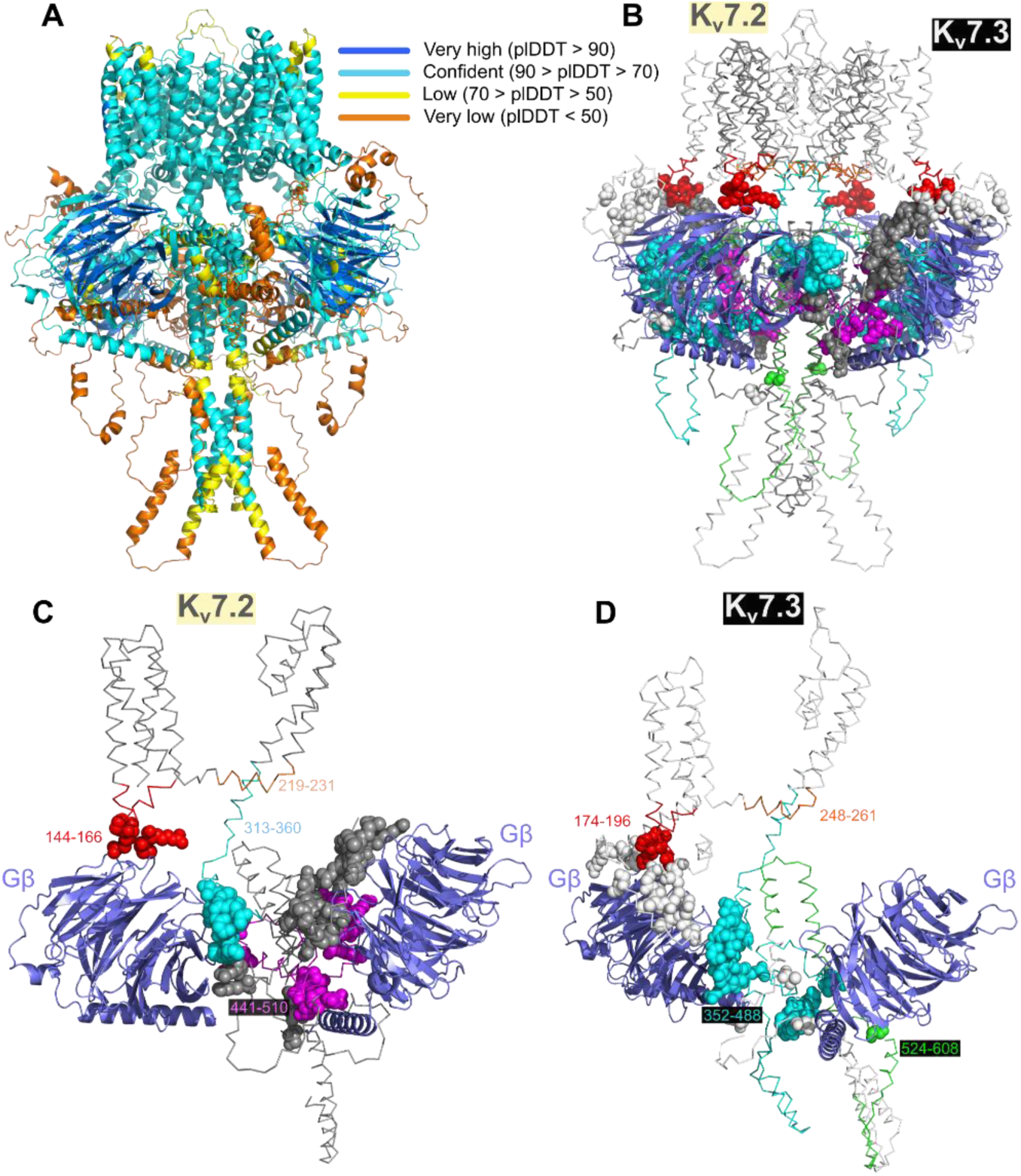
AlphaFold-Multimer model reveals candidate Gβγ-binding regions in the K_V_7.2/7.3 channel. **(A)** Full assembly colored by per-atom confidence estimate. **(B)** Same model, with channel shown as ribbon, K_V_7.2 in off-white, K_V_7.3 in gray, and Gβ in purple. The peptide array-mapped regions of K_V_7.2 and K_V_7.3 are colored as in figure 5: K_V_7.2 residues 144-166 (S2-S3), 219-231 (S4-S5 linker), 313-360 (proximal C-terminus/helix A), and 441-510; K_V_7.3 residues 174-196, 248-261, 352-488, and 524-608 (helix B-C). Residues interacting with Gβ are shown as spheres. There is an overlap between the peptide array-mapped regions and the AlphaFold model. **(C and D)** Close-ups of the cytoplasmic interface for K_V_7.2 **(C)** and K_V_7.3 **(D)** showing candidate segments converging toward Gβ.

**Fig. S8.**
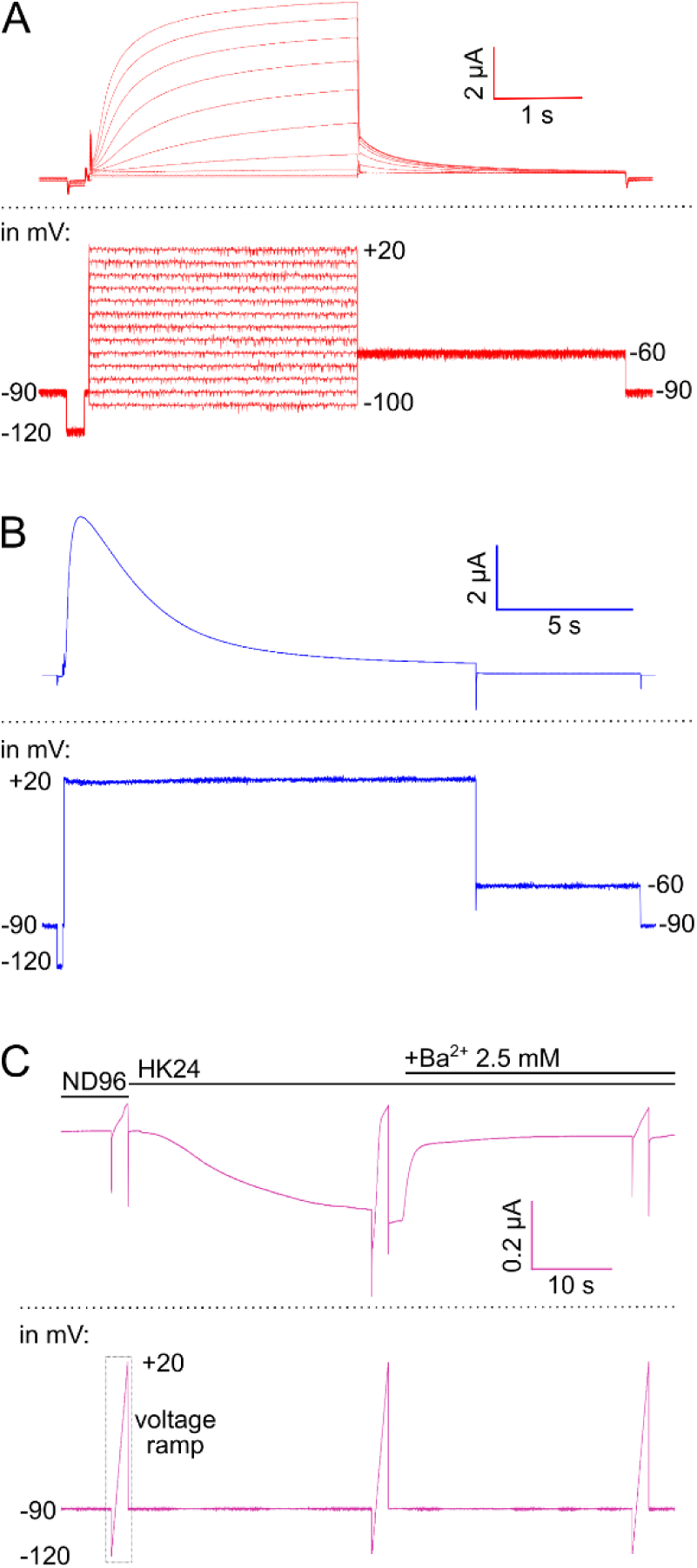
Examples of recording protocols (bottom panels) and representative current records (upper panels). **(A)** Voltage step protocol sequence (for I-V relationships): -90 mV (200 ms) → - 120 mV (200 ms) → -90 mV (50 ms) → -100 mV to +20 mV in 10 mV increments (3000 ms) → - 60 mV (3000 ms) → -90 mV (200 ms). **(B)** PIP_2_ depletion protocol sequence: -90 mV (200 ms) → -120 mV (200 ms) → -90 mV (50 ms) → +20 mV (15000 ms) → -60 mV (6000 ms) → -90 mV (200 ms). **(C)** For the recording of GIRK currents oocytes were held at -80 mV throughout the whole recording and only solutions were replaced: ND96 (5 s) → Voltage ramp from -120 to 50 mV (2 s) → HK24 solution (in mM: 24 KCl, 74 NaCl, 1 MgCl2, 1 CaCl2, 5 HEPES, pH adjusted to 7.5 with KOH; 30 s) → Voltage ramp from -120 mV to 50 mV (2 s) → HK24 + GIRK blocker (2.5 mM Ba^2+^).

### Supplementary Tables

**Table S1:**
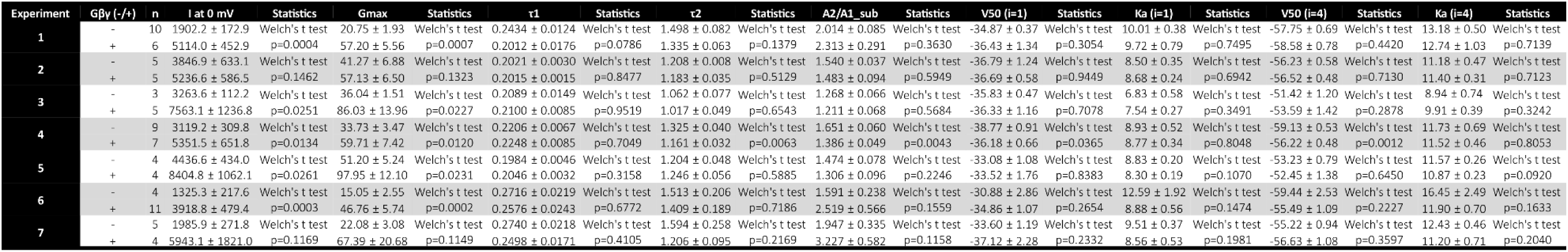

**Table S2:**
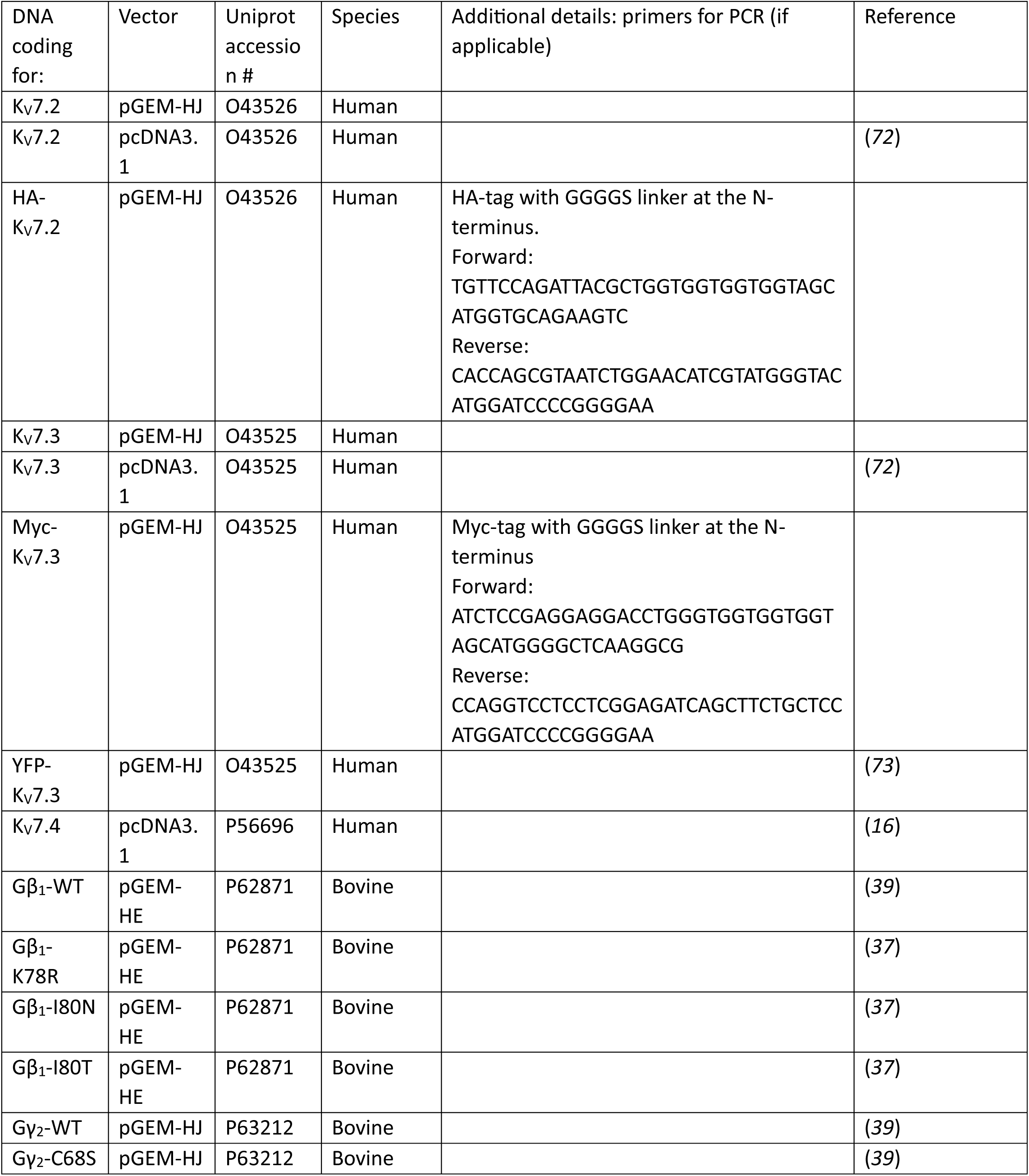
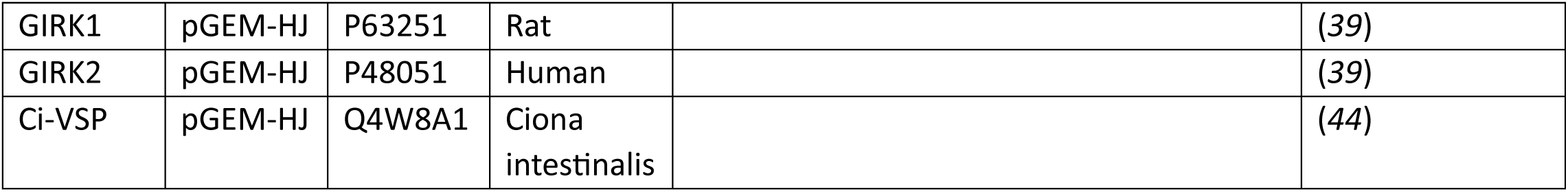
DNA constructs used in this study.

**Table S3:** antibodies used in this study.

| Name | Catalog # | RRID | Host | Target |
| --- | --- | --- | --- | --- |
| ChromPure® Donkey IgG, whole molecule | 017-000-003 | AB_2337256 | Donkey |  |
| GNB1 antibody | GTX114442 | AB_10619473 | rabbit | Human, Mouse, Rat, Zebrafish |
| Anti-GFP antibody | ab6556 | AB_305564 | rabbit |  |
| HA-probe Antibody (F-7) Alexa Fluor® 647 | sc-7392 AF647 | AB_627809 | Mouse |  |
| Anti-Myc/c-Myc Antibody (9E10) Alexa Fluor® 488 | sc-40 AF488 | AB_2892598 | mouse |  |
| Cy™3 AffiniPure® Donkey Anti-Rabbit IgG (H+L) | 711-165-152 | AB_2307443 | Donkey | Rabbit |
| Anti-KCNQ2 antibody | GTX82891 |  | Rabbit | Human, Mouse, Rat |
| Anti-KCNQ3 antibody | APC-051 | AB_2040103 | Rabbit | Human, Mouse, Rat |
| Anti-KCNQ4 antibody | APC-164 | AB_2341042 | Rabbit | Human, Mouse, Rat |
| Anti-Gβ (H-1) antibody | Sc-166123 |  | Mouse | Human, Rat |
| Duolink PLA probe Anti-rabbit minus | DUO82005 |  | Donkey |  |
| Duolink PLA probe Anti-mouse plus | DUO82001 |  | Donkey |  |

